# Structural Diversity and Dynamics of Metabotropic Glutamate Receptor/Beta-Arrestin Coupling

**DOI:** 10.1101/2025.02.03.636340

**Authors:** Dagan C. Marx, Kevin Huynh, Alberto J. Gonzalez-Hernandez, Alexa Strauss, Carlos Rico, Dirk Siepe, Pamela N. Gallo, Joon Lee, Sheida Sharghi Moshtaghin, Anisul Arefin, Willem F. Weber, Johannes Broichhagen, David Eliezer, Marian Kalocsay, George Khelashvili, Joshua Levitz

## Abstract

Beta-arrestins (β-arrs) are cytosolic proteins which mediate G protein-coupled receptor (GPCR) desensitization, endocytosis, and signaling. Despite the widespread physiological roles of β-arr coupling, the molecular basis of GPCR/β-arr interaction has been studied primarily in monomeric family A GPCRs. Here we take an integrative biophysical and structural approach to uncover molecular diversity in β-arr coupling to the neuromodulatory metabotropic glutamate receptors (mGluRs), prototypical, dimeric family C GPCRs. We find, using a new single molecule pulldown assay, that mGluRs couple to β-arrs with a 2:1 or 2:2 stoichiometry via a combination of “tail” and “core” interactions. Using single molecule FRET analysis, we also find that β-arr1 stabilizes active conformations of mGluR8. Cryo-EM structures of mGluR8 alone or with either G proteins or β-arr1 reveal transducer-specific mGluR8 active states and, in combination with molecular dynamics simulations, define the positioning of mGluR8-bound β-arr1, supporting a steric mechanism of mGluR desensitization involving interactions with both subunits and the lipid bilayer. Finally, combinatorial mutagenesis enables the identification of a landscape of homo- and hetero-dimeric mGluR/β-arr complexes, including mGluR/β-arr1/β-arr2 megacomplexes, providing a framework for family C GPCR/β-arr coupling and expanding the known range of GPCR/transducer coupling modes.

## Introduction

G protein-coupled receptors (GPCRs) are finely tuned transmembrane signaling molecules that sense extracellular stimuli and activate a wide range of intracellular effector pathways^1^. The precise signaling of GPCRs is central to diverse physiological processes, underscoring their role as major drug targets^2, 3^. Given their biological and therapeutic significance, great effort has been undertaken to understand the regulatory processes that control GPCR function. A major form of GPCR regulation depends on the multifunctional, cytosolic β-arrestins (β-arr1, β-arr2) which bind to activated GPCRs that have been phosphorylated by G protein-coupled receptor kinases (GRKs)^4, 5^. Following recruitment and complex formation, β-arrs can desensitize GPCR signaling by sterically blocking G protein access and initiating receptor internalization via clathrin recruitment. β-arr mediated receptor internalization can be subdivided into two classes: receptors that transiently recruit β-arrs to the plasma membrane but do not co-internalize and typically undergo endosomal recycling (class A), and receptors that form stable β-arr complexes that persist into endosomes and typically undergo lysosomal degradation (class B)^6–10^. In addition to their roles in receptor desensitization, β-arrs can also serve as scaffolds to initiate secondary waves of signaling^11^.

Biophysical and structural studies have shown how phosphorylated GPCR C-terminal tails mediate β-arr recruitment and activation^12–17^ and how β-arrs undergo stepwise recruitment via “tail” and “core” interactions with GPCRs^6, 18–23^. However, the details of β-arr coupling can vary dramatically between GPCR subtypes^6, 24–32^, between β-arr subtypes^24, 33–36^, and between different ligands for the same GPCR^37–43^, motivating ongoing studies of β-arrs across the vast landscape of GPCR signaling and regulation.

β-arr coupling has primarily been studied in the largely monomeric family A and family B GPCRs, leaving potential differences in other branches of the diverse GPCR superfamily uncharacterized. Family C GPCRs, including the prototypical metabotropic glutamate receptors (mGluRs), the GABA_B_ receptors (GABA_B_R), and calcium-sensing receptor (CaSR), possess distinctive structural features, including large extracellular ligand binding domains (LBDs) and constitutive dimerization^44, 45^. These unique properties raise key questions about activation and coupling to G protein and β-arr transducers, including: what stoichiometries do receptor/transducer complexes adopt in homo- or heterodimeric GPCRs? What are the structural and conformational determinants of receptor/transducer coupling?

Structural studies of G protein-bound family C GPCRs have begun to provide a framework for understanding G protein-coupling and have revealed a strict 2:1 GPCR: G protein heterotrimer stoichiometry^46–53^. Spectroscopic studies have confirmed that G proteins stabilize active agonist-driven receptor conformations^54–57^. However, family C GPCR/G protein structures do not show canonical signatures of transmembrane domain (TMD) activation, including outward motions of TM6, as observed in monomeric GPCRs^58–61^. While far less is known about family C GPCR/β-arr coupling, we and others have shown recently that a subset of mGluRs, including mGluR3, mGluR7, and mGluR8, undergo GRK and β-arr dependent desensitization, endocytosis, and subtype-specific intracellular trafficking fates^62–66^. mGluR heterodimerization further diversifies this family^67–69^, enabling β-arr-dependent endocytosis of otherwise-resistant subtypes (e.g. mGluR2) and altering receptor trafficking fate (i.e. endocytic recycling versus lysosomal degradation)^63^. Differences in β-arr-dependent internalization and trafficking fates have largely been attributed to distinct patterns of phosphorylatable residues in the receptor C-termini^62, 63, 70^, although direct analysis of mGluR/β-arr complexes is lacking, limiting our molecular understanding of mGluR/β-arr coupling.

Given the central role of mGluRs in various modes of neuromodulation and their potential as drug targets for neurological and psychiatric disorders^71, 72^, a comprehensive biophysical and structural picture of mGluR coupling to β-arrs is needed. In this study, we develop a single molecule pulldown (SiMPull) imaging assay to directly assess GPCR/β-arr complex formation and characterize β-arr coupling across the mGluR family. We find variability in coupling propensity across subtypes, including the formation of both 2:1 and 2:2 mGluR/β-arr complexes. Mass spectrometry and mutational scanning, along with single molecule Förster resonance energy transfer (smFRET) and negative stain electron microscopy (EM), produce a mechanistic picture of “tail” and “core” interactions which drive mGluR8/β-arr coupling. Cryo-EM structures of mGluR8, mGluR8/G protein, and mGluR8/β-arr reveal subtle differences in transducer-specific active state TMD conformations. Guided by cryo-EM structures, molecular dynamics simulations and mutagenesis analysis define dynamic β-arr1 positioning in 2:1 core-bound complexes. Finally, structure-guided SiMPull analysis reveal a range of 2:1 and 2:2 mGluR8/β-arr binding modes, including formation of cis and trans-coupling between CTDs and TMD cores, heterodimeric mGluR8/mGluR2/β-arr complexes, and mGluR8/β-arr1/β-arr2 megacomplexes, expanding the diversity of known GPCR/β-arr assemblies.

## Results

### A single molecule assay to assess GPCR/β-arr1 complex formation

To directly detect and quantitate mGluR/β-arr complexes, we sought to establish a single molecule imaging platform based on visualizing constructs bearing self-labeling tags (e.g. SNAP, Halo) for organic fluorophore conjugation. We turned to previously established N-terminally SNAP-tagged mGluR constructs^67^ and generated a C-terminally Halo-tagged β-arr1 (*see Methods*). Consistent with our prior work^62, 63^, live cell imaging revealed that SNAP-mGluR3 and SNAP-mGluR8 (labeled with cell impermeable SBG-JF_646_) recruit cytosolic β-arr1-Halo (labeled with cell permeable CA-JF_549_ for Halo) to the plasma membrane upon glutamate addition (**Fig. 1A, B; Fig. S1A-D**). β-arr1-Halo co-internalized with SNAP-mGluR8, but not with SNAP-mGluR3 (**Fig. S1A-D**), further confirming that the unique subtype-specific properties of β-arr recruitment are maintained with our constructs. We also prepared a C-terminally truncated β-arr1 construct (“β-arr1-ΔCTD-Halo”) where the autoinhibitory C-terminal domain (aa 383-418) of β-arr1 has been removed. This construct was efficiently recruited to the plasma membrane by both mGluR3 and mGluR8 (**Fig. S1E, F**) and impaired internalization of both receptor subtypes, consistent with the removal of motifs which facilitate clathrin binding within the β-arr1-CTD (**Fig. S1G, H**)^73–77^.

**Figure 1.**
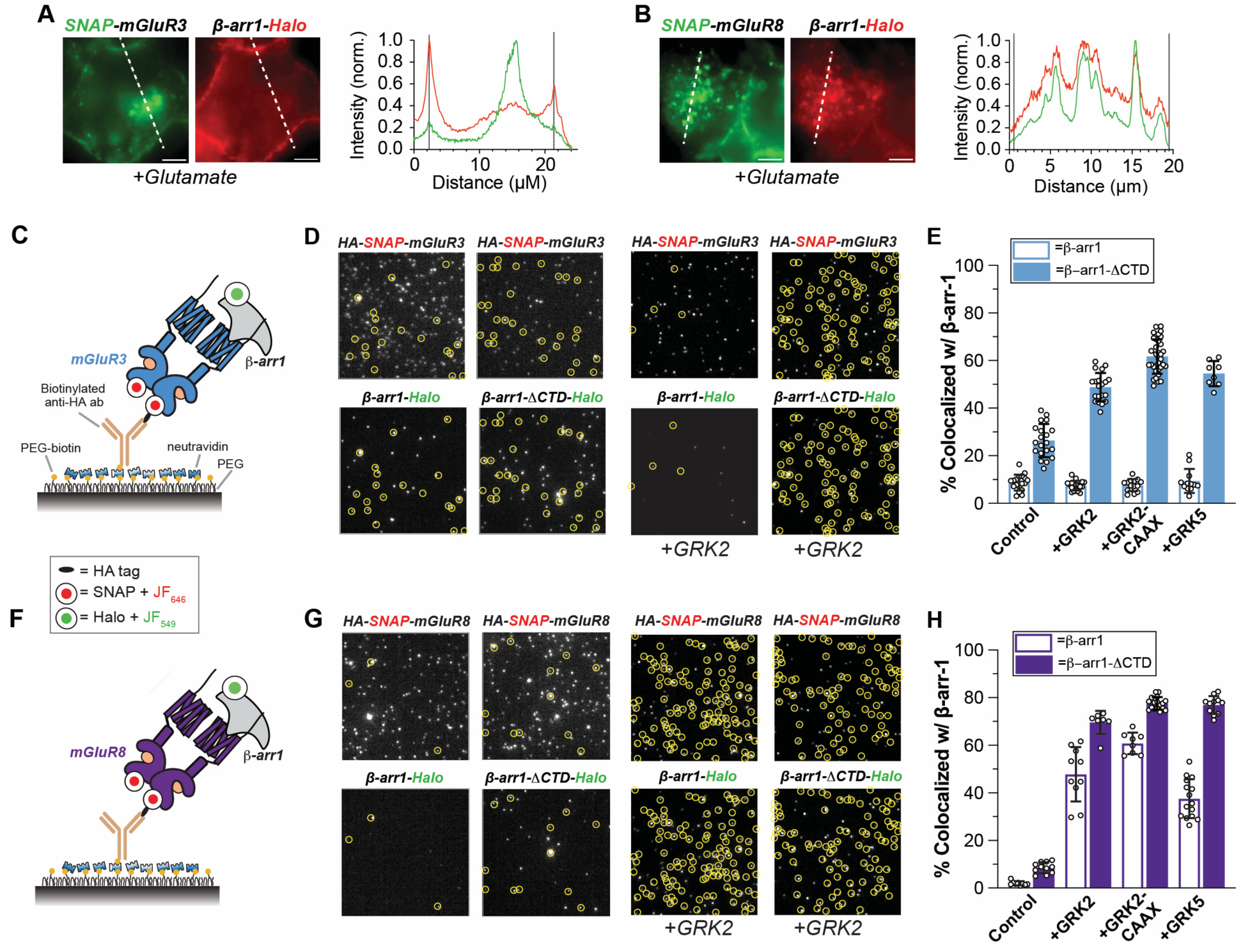
A single molecule assay for detection of mGluR/β-arr complexes. **(A-B)** Representative images showing surface localization and co-internalization of β-arr1 with mGluR3 (A) or mGluR8 (B), respectively. **(C-H)** Schematic (C, F), representative images (D, G), and summary quantification (E, H) of SiMPull assay of mGluR/β-arr1 complexes. Yellow circles (D, G) indicate colocalized spots indicative of mGluR/β-arr1 complexes. Points represent individual movies (E, H). All data shown as mean ± SEM. Scale bar: 5 µm.

We then assessed the ability of SNAP-tagged mGluRs to couple to β-arr1 via single molecule pulldown (“SiMPull”)^78, 79^. Briefly, fresh lysates were prepared from cells transfected with SNAP-tagged mGluRs and Halo-tagged β-arr1 constructs that were labeled with fluorophores and subsequently incubated with saturating concentrations of glutamate. An N-terminal HA-tag on the SNAP-mGluR constructs enabled receptor isolation on passivated glass coverslips via a conjugated anti-HA primary antibody (**Fig. 1C-H**). We then used total internal reflection fluorescence (TIRF) microscopy to visualize individual receptors and β-arr1 in separate channels and to quantify complex formation via colocalization of spots in both channels.

In the absence of GRK overexpression, minimal complex formation (<10%) with either WT or β-arr1-ΔCTD was observed for mGluR8, while mGluR3 showed modest colocalization with β-arr1-ΔCTD (∼25%) (**Fig. 1D-H**). Co-transfection with either GRK2, GRK5, or a membrane-tethered GRK2 containing a C-terminal CaaX-box motif (GRK2-CaaX), dramatically increased the extent of β-arr1 colocalization for both mGluR3 and mGluR8 (**Fig. 1D-H)**, likely by enhancing the degree of receptor phosphorylation. Both mGluR3 and mGluR8 produced substantial pulldown of β-arr1-ΔCTD with ∼60% or 75% of receptors, respectively, showing β-arr1 colocalization under optimal conditions with GRK2-CaaX co-transfected. Strikingly, only mGluR8 showed substantial co-localization (∼55%) with WT β-arr1 (**Fig. 1D-H**). The strong dependence on β-arr1 truncation for colocalization with mGluR3, but not mGluR8, was seen across all GRK conditions (**Fig. 1E, H**). Importantly, in the absence of receptor co-expression, very minimal β-arr1-Halo pulldown was observed (**Fig. S1G**). Overall, these results suggest that mGluR8 has a higher phosphorylation threshold for β-arr1 coupling but can form more stable complexes with β-arr1 than mGluR3, which may explain the ability of mGluR8/β-arr1 complexes to persist following endocytosis.

To assess the generalizability of our assay, we investigated β-arr1 coupling to two SNAP-tagged prototypical family A GPCRs, the beta-2 adrenergic receptor (β2AR) and the arginine vasopressin 2 receptor (V2R). In the presence of GRK2-CaaX and saturating agonist, each receptor showed pulldown of β-arr1-ΔCTD, but V2R was substantially more efficient (∼60% vs. 20% colocalization). Similarly to mGluR8 and mGluR3, respectively, V2R was able to produce strong pulldown of WT β-arr1 while β2AR was not (**Fig. 2A, B**). We confirmed that both receptors recruit β-arr1-Halo in live cells but SNAP-β2AR interacts transiently^7, 80^ while SNAP-V2R co-internalizes^7^ with with β-arr1-Halo (**Fig. S2A-D**). These results are consistent with class B (V2R, mGluR8), but not class A (β2AR, mGluR3), GPCRs being capable of stable interactions with WT β-arr1 and indicate that our SiMPull assay can provide a general means of quantitatively probing and classifying GPCR/β-arr interactions. We interpret the colocalization percentages of β-arr1 with receptors to report on the relative stability of the complex, but these *in vitro* percentages may be underestimated due to both the dissociation of some complexes upon cell lysis and incomplete fluorescent labeling (see Methods).

**Figure 2.**
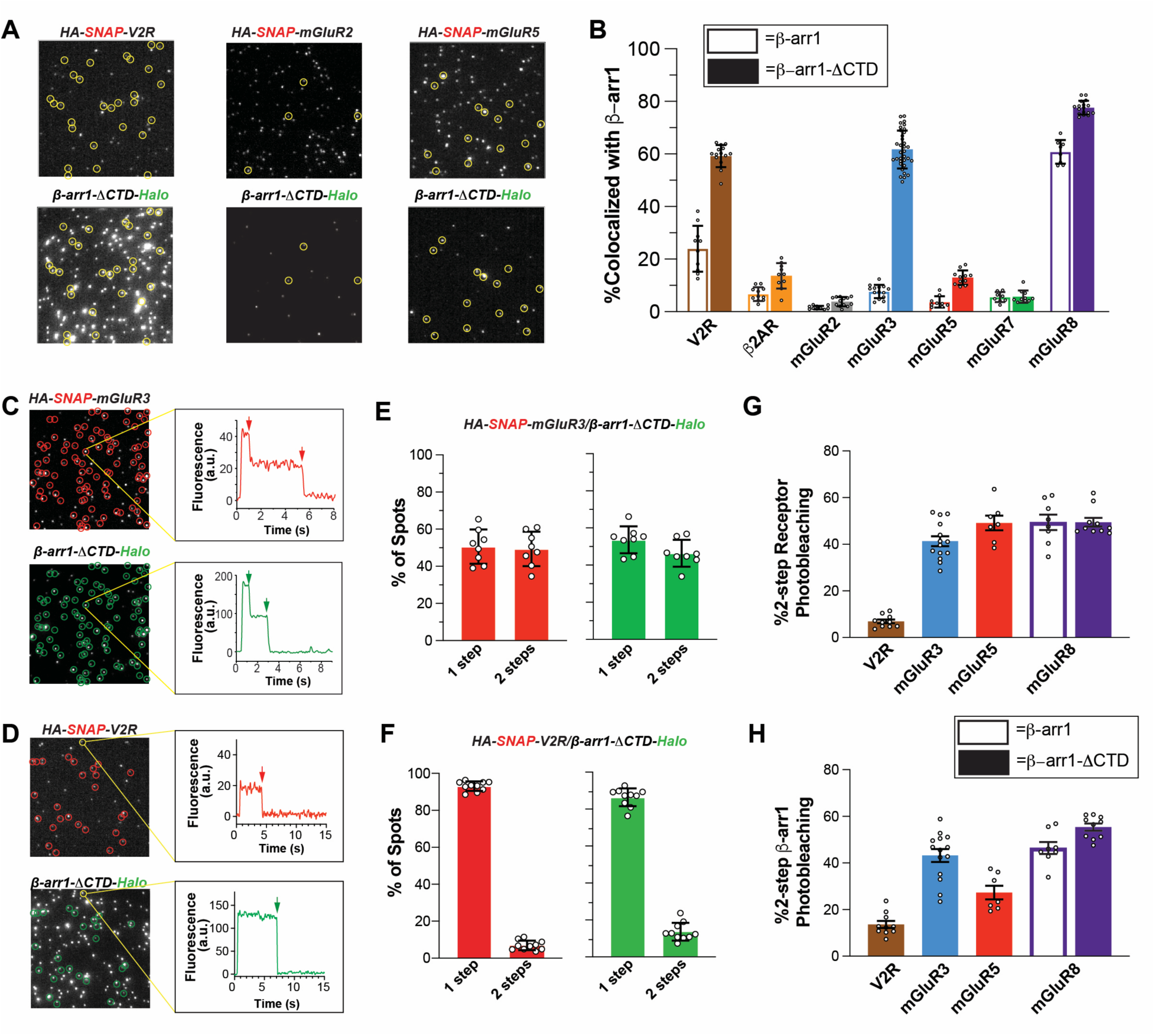
Single molecule analysis of GPCR/β-arr1 coupling reveals a range of efficiencies and stoichiometries, including 2:2 mGluR/β-arr1 complexes. **(A)** Representative images showing variable b-arr1-DCTD pulldown efficiencies for V2R, mGluR2, and mGluR5. **(B)** Summary graph of β-arr1 colocalization across GPCRs for both WT and β-arr1-DCTD. **(C-F)** Representative images and bleaching step traces **(C, D)** and photobleaching step distributions (E, F) for β-arr1-DCTD complexes mGluR3 and V2R. Arrows indicate bleaching steps. **(G-H)** Summary of % of colocalized molecules showing 2-step bleaching in the receptor (G) and β-arr1 (H) channels across receptors. For all experiments, GRK2-CaaX was co-expressed and cells were treated with agonist. Circles (A, C, D) indicate colocalized spots indicative of GPCR/b-arr1 complexes. Points represent individual movies (B, E-H). All data shown as mean ± SEM.

We next extended our analysis across the mGluR family and tested SNAP-tagged mGluR2, mGluR5, and mGluR7 in the presence of GRK2-CaaX and either β-arr1-Halo or β-arr1-ΔCTD-Halo (**Fig. 2A, B**). SNAP-mGluR2 showed negligible levels (<5%) of co-localization with either β-arr1 or β-arr1-ΔCTD (**Fig. 2A, B**), consistent with the lack of β-arr1 recruitment and receptor internalization observed in live cells (**Fig. S2E, F**)^62, 63^. In contrast, SNAP-mGluR5 showed modest pulldown (∼15-20%) of β-arr1-ΔCTD, but not WT β-arr1 (**Fig. 2A, B**). Live-cell imaging mirrored these results where only β-arr1-ΔCTD, showed detectable glutamate-induced membrane localization (**Fig. S2G, H**). Finally, SNAP-mGluR7 showed weak (∼5-10%) pulldown of both β-arr1-ΔCTD, and WT β-arr (**Fig. 2B**), in line with live cell imaging showing modest recruitment of β-arr1-ΔCTD, but not WT β-arr1 (**Fig. S2I, J**).

### Variable stoichiometries of GPCR/β-arr1 complexes

A major advantage of single molecule imaging is the ability to measure the stoichiometry of protein assemblies via photobleaching step analysis, a technique we have used to confirm the constitutive dimerization of mGluRs^68, 69, 81^. We thus asked two key questions about mGluR/β-arr1 complexes: 1) Do mGluRs retain dimerization upon β-arr interaction? And 2) how many β-arr1 molecules can interact with an mGluR at a time? To assess these questions, we analyzed the photobleaching step distribution for both the receptor and β-arr1 channels for all β-arr-colocalized mGluR3, mGluR5, and mGluR8 spots (**Fig. 2C-H**). We found that ∼50% of spots displayed 2 bleaching steps in the receptor channel for each mGluR subtype (**Fig. 2G**), confirming that mGluRs remain dimeric upon β-arr interaction. This result is in line with our recent study showing that internalized mGluR3 and mGluR8 are strict dimers^63^. As previously established^68, 69, 81^, the subpopulation of mGluRs showing single bleaching steps is an expected phenomenon for a strictly dimeric protein complex due to variety of factors including incomplete fluorophore labeling and incidental bleaching.

Surprisingly, ∼25-55% of co-localized complexes also showed 2 bleaching steps in the β-arr1 channel (**Fig. 2C, E, H; Fig. S2K**), indicating that a combination of 2:1 and 2:2 mGluR:β**-**arr1 complexes can form. The proportion of 2-step β-arr1 bleaching steps was lower for mGluR5 than mGluR3 or mGluR8 which may be due to lower overall complex formation for this receptor (**Fig. 2H**). As a key control, V2R:β-arr1-ΔCTD complexes showed background levels (∼10%) of 2-step bleaching in both channels (**Fig. 2D, F-H**) consistent with formation of a 1:1 complex between monomeric family A GPCRs and a single β-arr1 molecule. To further validate our finding of 2:2 mGluR:β-arr1 complexes we sought to directly image β-arr1 colocalization at a single receptor. To do this, we labeled β-arr1-Halo with a mix of both CA-JF_549_ and CA-JF_646_ fluorophores, while leaving the SNAP-tag on mGluR8 unlabeled (**Fig. S3A**). We observed substantial colocalization (∼40%) of β-arr1-ΔCTD when imaging both channels (**Fig. S3B**), in agreement with our photobleaching step analysis implying the existence of mGluR/β-arr1 complexes containing two β-arr1 molecules bound to one receptor dimer.

### mGluR8 phosphorylation controls the strength of β-arr1 interaction

Having observed particularly strong colocalization of mGluR8 and β-arr1, we decided to focus our analyses on better understanding this complex. Our prior work in cells has pointed to a critical role of phosphorylation sites in the mGluR8 C-terminus^63^ for β**-**arr mediated receptor internalization. Mass spectrometry of purified, agonist-treated mGluR8 showed extensive phosphorylation in the distal serine/threonine (S/T)-rich region (aa888-907), including many double-phosphorylated peptides (**Fig. S4**). This finding is consistent with the likelihood that multiple phosphorylation sites simultaneously contribute to β-arr recruitment^12, 14, 82, 83^.

To understand the role of each putative mGluR8 phosphosite on β**-**arr1 complex formation, we performed alanine scanning mutagenesis of the 11 potential phospho-sites in the mGluR8 S/T-rich region. We tested each mutant in SiMPull using our highest colocalization condition (GRK2-CaaX, β-arr1-βCTD-Halo) and a less tightly coupled condition (GRK2, β-arr1-Halo), finding similar trends in both conditions with larger effects for the latter (**Fig. 3A**). We found no pulldown with simultaneous mutation of all 11 phospho-sites (“mGluR8-11xA”), highlighting the need for phosphorylation in the mGluR8 S/T-rich region for β-arr1 coupling. All individual mutants showed impairment with the largest effects seen for T896A, T898A, and T899A, residues that could constitute a recently identified arrestin binding PxPP motif^14, 84^. Consistent with this, a triple mutation of T896A/T899A/T898A (“mGluR8-3xA”) decreased pulldown to the same degree as mGluR8-11xA, suggesting a dominant role for this PxPP motif in β-arr1 coupling (**Fig. 3A**).

**Fig. 3.**
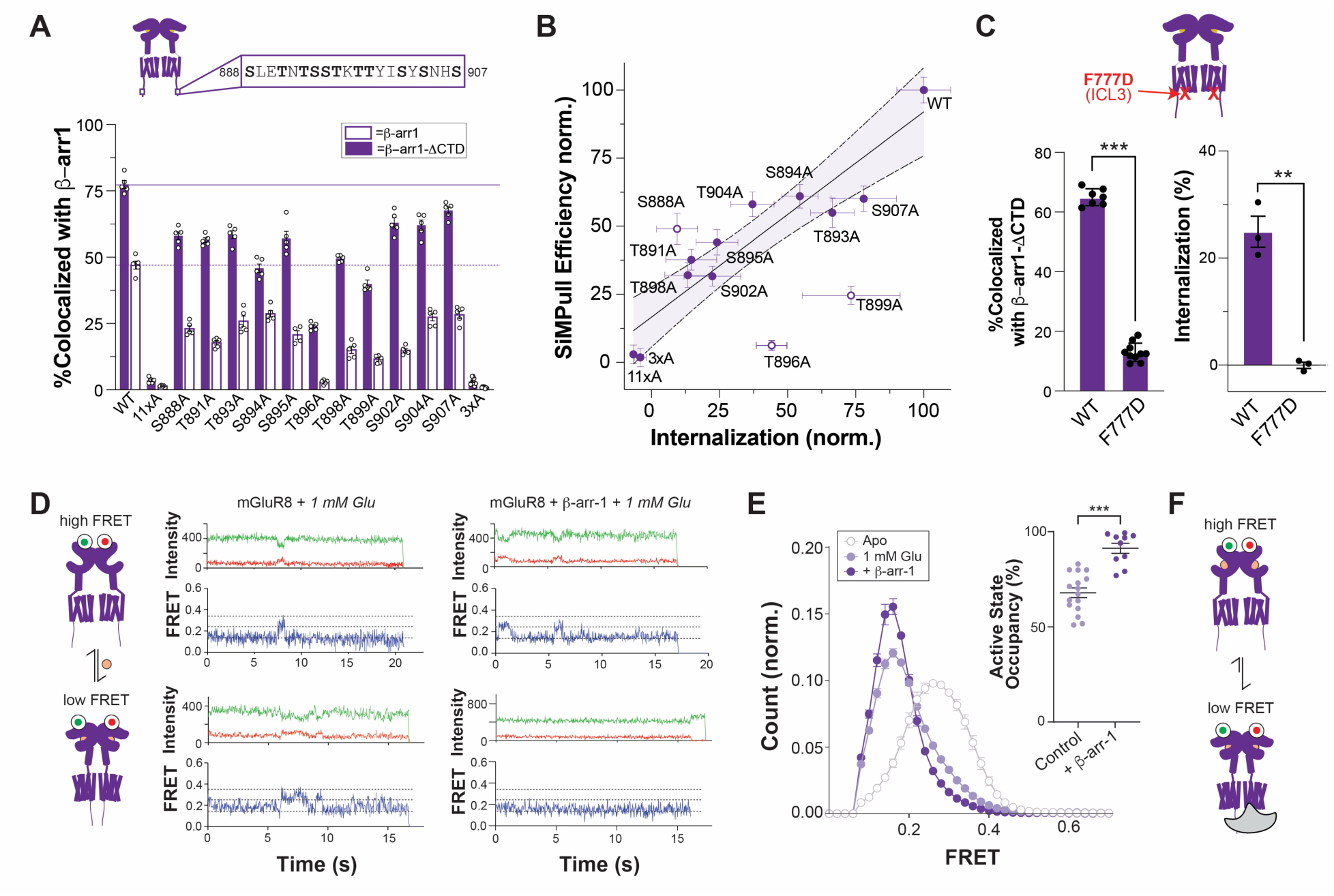
Defining tail and core interactions that drive mGluR8/β-arr1 coupling. **(A)** Summary graph of SiMPull data for colocalization of all mGluR CTD point mutations either with wild type β-arr1-Halo (with GRK2 co-expression) or β-arr1-βCTD (with GRK2-CaaX co-expression). 3xA=T896A/T898A/T899A. **(B)** Scatter plot showing correlation between effects of mutations on β-arr1 co-localization in SiMPull (y-axis) and internalization in cell-based surface labeling assay (x-axis). A linear fit to the data excluding outliers (S888A, T896A, T899A) is shown with the 95% confidence interval shaded. **(C)** Summary graphs showing impairment of β-arr1βCTD colocalization in SiMPull (left) and mGluR8 internalization (right) with the F777D mutation in ICL3. **(D)** Left, schematic showing inter-LBD FRET assay where glutamate binding decreases FRET levels due to LBD closure and reorientation. Right, representative smFRET traces showing occasional transitions between low, medium, and high FRET states (dotted lines). **(E)** Histograms showing FRET distributions in the absence or presence of 1 mM Glutamate. β-arr1 increases the height of the low FRET peak. Inset shows occupancy of the low FRET 0.15 peak as calculated from Gaussian fits across individual experiments. **(F)** Summary schematic showing that β-arr1 stabilizes the active, low FRET conformation of mGluR8. Points represent individual movies (A, C, E) or separate experimental days (C, right). Unpaired t-test is used in (C, E). **, p<0.01; ***, p<0.001. All data shown as mean ± SEM.

We then assessed the effects of each CTD variant using an established live cell imaging-based assay of glutamate-induced mGluR8 internalization^62, 63^ to understand how differential phosphorylation impacts β-arr1-mediated receptor desensitization. We observed clear impairment of receptor internalization for most mutations (**Fig. S5A, B**), including a complete loss of internalization for both mGluR8-11xA and mGluR8-3xA. When comparing our colocalization and internalization datasets for each mGluR8 CTD mutant, we found a mostly linear correlation between the extent of impairment of β-arr1 colocalization and the extent of impairment of ligand-induced internalization (**Fig. 3B**). However, mutations to three phospho-sites had differing effects in our two assays: S888, T896, T899. S888A impacted β-arr1 SiMPull colocalization less profoundly than internalization, while T896A and T899A impaired binding more than internalization (**Fig. 3B**).

To better understand the discrepancy between our two assays for these three variants, we directly imaged β-arr1 recruitment and receptor endocytosis in live cells. Surprisingly, S888A revealed an almost complete loss of β-arr1 recruitment and receptor internalization (**Fig. S5C**). One possible explanation is that S888 serves as a gatekeeper residue whose phosphorylation helps drive subsequent phosphorylation of other positions. This could explain why SiMPull effects under conditions where GRK2-CAAX is overexpressed are less pronounced. In contrast, T899A showed clear receptor internalization and β-arr1 recruitment to the plasma membrane, but minimal co-localization of β-arr1 with internalized receptors (**Fig. S5D**). A potential explanation is that phosphorylation of T899 is critical for maintaining a conformation of β-arr1 that survives endocytosis but is not required for initial β-arr1 recruitment to the plasma membrane. Similarly, T896A showed impaired recruitment of β-arr1 but still maintained some weak endosomal co-localization (**Fig. S5E**). Motivated by these complex phenotypes, we tested glutamate-induced degradation using our established SNAP-tag based assay^63^ and found that T896A, T898A, and T899A all showed larger impairments compared to their effects on receptor internalization (**Fig. S5F**). This finding is consistent with the interpretation that stable β-arr coupling is more critical for complex co-internalization and degradation than internalization. Taken together, the myriad effects of mutations across the ST-rich region of the mGluR8-CTD, suggest the potential of multiple β-arr1 binding modes.

### mGluR8/β-arr1 core coupling stabilizes active mGluR8 conformations

We next asked if the mGluR8 TMD core contributes to mGluR8/β-arr1 coupling. We first turned to a well-established mutation of a highly conserved phenylalanine (F777 in mGluR8) in intracellular loop 3 that strongly impairs G protein coupling across mGluR subtypes^69, 85, 86^. We hypothesized that if a similar interface is used for β-arr1 core coupling, mutation to this position would also strongly impair mGluR8/β-arr1 coupling and endocytosis^62^. Colocalization of β-arr1 was strongly reduced in SiMPull and glutamate-driven internalization in live cells was abolished for mGluR8-F777D (**Fig. 3C; Fig. S6A, B**). While such effects could be due, at least in part, to impairment of GRK coupling, this result supports the hypothesis that TMD core interactions contribute to mGluR8/β-arr1 coupling. To further test this hypothesis, we asked if receptor core interactions alone are sufficient to produce detectable mGluR8/β-arr1 coupling in cells by producing an mGluR8 construct lacking a CTD (mGluR8-ΔCTD). In line with a role for core coupling, SNAP-mGluR8-βCTD showed weak, but clear, glutamate-driven internalization (**Fig. S6C, D**).

As we hypothesize that mGluR8/β-arr1 coupling involves interactions with the TMD core, we asked if β-arr1 coupling alters the conformational dynamics of mGluR8. Prior work using cryo-EM and FRET-based assays has established that G proteins can stabilize mGluRs in their active-like conformation^46–48, 54–56^, but whether β-arrs exert a similar or, potentially, opposite effect as part of an inhibitory mechanism, is not clear. We tested this using an inter-LBD single molecule (smFRET) assay that has been used to characterize agonist-induced transitions from inactive, high FRET, to active, low FRET, conformations^54, 56, 87–90^ (**Fig. 3D**). Application of saturating glutamate shifted FRET histogram peaks of SNAP-mGluR8 immobilized from lysates of HEK293T cells from a center of ∼0.35 to a center of ∼0.15 but with a clear shoulder indicating incomplete occupancy of the active, low-FRET state (**Fig. 3E**). In both the absence (**Fig. S7A**) and presence of glutamate (**Fig. 3D**), a sub-population of molecules showed dynamic transitions between FRET states. This agrees with prior smFRET studies of group II/III mGluRs which showed variable degrees of incomplete occupancy of the low FRET state in response to orthosteric agonists^87, 88, 91^. When SNAP-mGluR8 lysates were prepared from cells co-expressing GRK2 and β-arr1- βCTD, the occupancy of the low FRET state in the presence of glutamate was substantially enhanced (**Fig. 3E**). This was accompanied by a reduction in the percentage of molecules showing dynamic transitions (**Fig. 3D; Fig. S7B**), suggesting that β-arr1 helps to “lock” mGluR8 in an active conformation. Hidden Markov modeling of high (0.35), medium (0.25), and low (0.15) FRET states revealed an increase in the dwell time in the low FRET state, but not the medium or high FRET states (**Fig. S7C**), in the presence of β-arr1, likely owing to decreased transitions between the low FRET active state and medium FRET state (**Fig. S7D**). Together these data show that β-arr1 promotes an active mGluR8 conformation by reducing transitions to inactive states (**Fig. 3F**), suggesting that β-arr1 recognizes a similar active state to G protein. Recent work has shown similar conformational effects of positive allosteric modulators (PAMs)^37, 56, 57, 92, 93^, indicating that they likely stabilize a similar active state or active states as β-arr1. Consistent with this, PAMs enhance both agonist-driven G protein and β-arr coupling for mGluR2, mGluR3, mGluR7, and mGluR8^37^.

### Negative stain EM reveals molecular diversity of mGluR8/β-arr1 complexes

To gain further insight into mGluR8/β-arr1 complexes, we first purified either full-length mGluR8 alone or mGluR8/β-arr1βCTD complexes in the presence of agonist (L-AP-4) and PAM (VU6005649; “VU600”). We turned to negative stain EM, which has been used to observe the presence of both “tail” and “core” bound β-arr1 for other GPCRs^14, 17, 24, 94^. Micrographs of the mGluR8 alone sample yielded 2-D classes showing dimeric mGluR8 with clearly resolved LBDs and TMD-containing detergent micelles in an orientation consistent with an active-like conformation of a mGluR (**Fig. 4A**). The mGluR8/β-arr1 complex micrographs contained particles with additional density on the intracellular side of the micelle consistent with mGluR8/β-arr1 complex formation (**Fig. S8**). Importantly, we observed no major change in mGluR8 LBD or CRD conformation when bound to β-arr1, confirming our smFRET-based finding that β-arr1 stabilizes an active state of mGluR8.

**Fig. 4.**
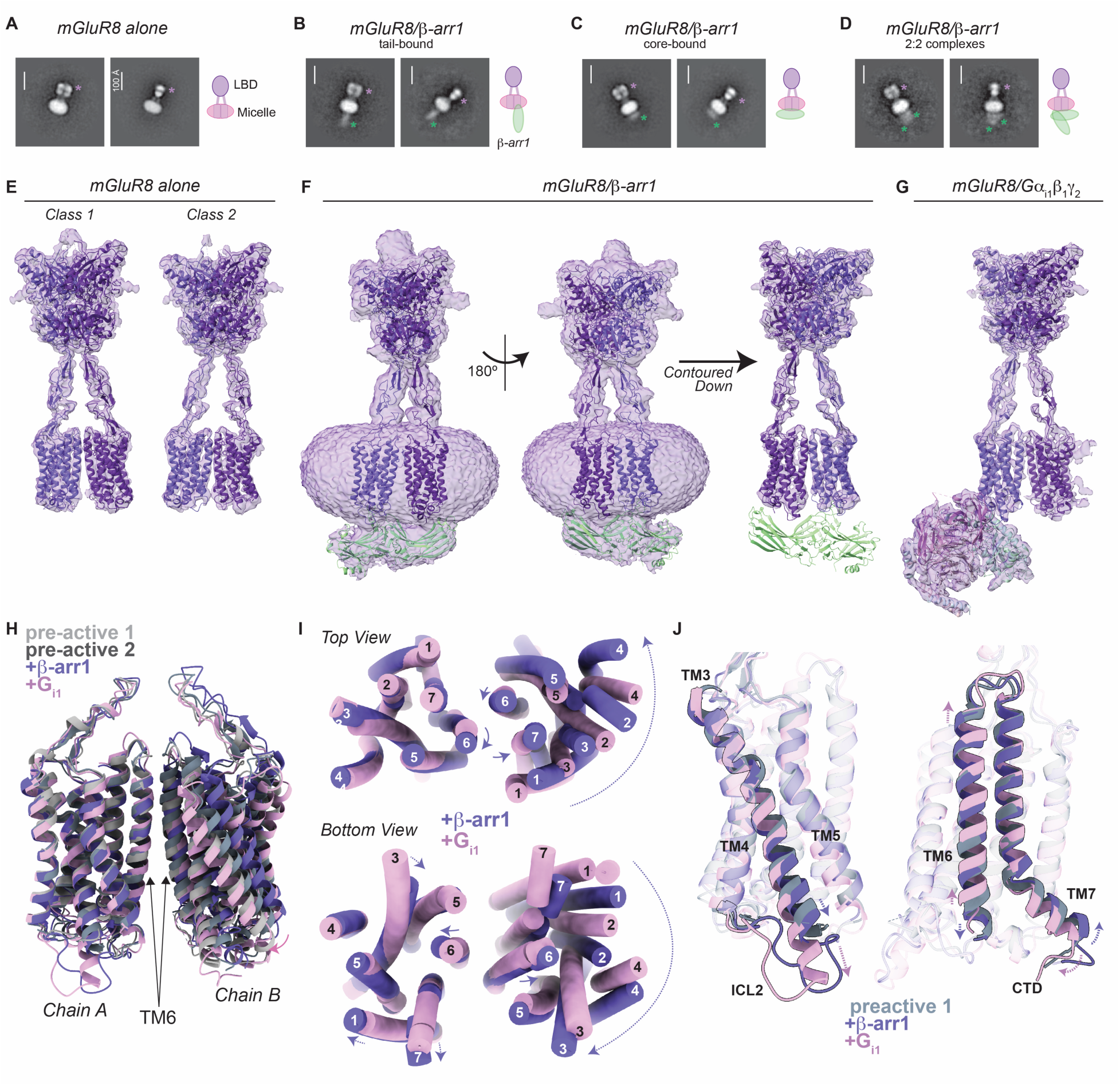
Structural analysis of agonist- and PAM-bound mGluR8, mGluR8/β-arr1, and mGluR8/G protein complexes. **(A-D)** Representative 2-D classes from negative stain analysis of mGluR8 alone (A) or mGluR8/β-arr1 (B) samples. A cartoon representation showing proposed ligand binding domain (LBD), micelle, and β-arr1 organization for each complex. **(E)** Full-length cryo-EM map and model of pre-active mGluR8 class 1 and class 2 structures bound to the orthosteric agonist L-AP-4 (green) and PAM VU600 (blue). **(F)** Multiple views showing mGluR8/β-arr1 cryo-EM map contoured to two different levels with β-arr1 aligned into the β-arr1 density. **(G)** mGluR8/G protein cryo-EM map and model. **(H)** TMD alignment **(I)** Top, extracellular view and, bottom, intracellular view, showing subtle rearrangement TMD dimer orientation between mGluR8/β-arr1 and mGluR8/G protein. Alignment is to the chain A TMD (left) in both (H) and (I). **(J)** Alignment of chain A TMD of mGluR8 structures highlighting repositioning, increased resolution and lengthening of the bottom of TM3 helix for transducer coupled structures (left), as well as subtle but opposite shifts of TM6 and TM7 in reference to the preactive 1 structure (right).

Focusing on the β-arr1 density, we identified multiple, distinct orientations of the β-arr1 relative to mGluR8. We were able to assign apparent “core” and “tail” bound binding modes based on the orientation, highlighting the conformational diversity of the mGluR8/β-arr1 complex (**Fig. 4B, C; Fig. S8**). Surprisingly, in the core-bound binding mode β-arr1 occupies a position that appears to be consistent with simultaneous interactions with both mGluR8 TMDs. Additionally, given our SiMPull evidence for 2:2 mGluR8/β-arr1 complexes, we manually inspected our micrographs and observed particles containing a larger density on the intracellular face of mGluR8, with a subset of these particles containing two separable densities (**Fig. S8**). 2-D classes generated from these particles revealed intracellular density that was larger than calculated for a single β-arr1 molecule and appeared to contain two distinct molecules with different orientations (**Fig. 4D**). In total, negative stain EM highlights the conformational and stoichiometric diversity of purified mGluR8/β-arr1 complexes that largely agrees with our single molecule fluorescence-based approaches.

### Cryo-EM structures of mGluR8, mGluR8/β-arr1, and mGluR8/G protein complexes

We next turned to cryo-EM to gain deeper insight into the structural basis of mGluR8/β-arr1 coupling. We determined the structure of pre-active mGluR8 bound to L-AP-4 and VU600 (**Fig. 4E; Fig. S9**), as there are no reported full-length transducer-free structures of mGluR8. Image processing revealed two classes with similar numbers of particles which each showed closed LBDs (**Fig. S10A**) with clear density for L-AP-4 in the inter-domain cleft of the LBDs (**Fig. S10B-D**) and active-like inter-LBD and inter-CRD orientations (**Fig. S10E, F**). The major difference between class 1 and class 2 structures is a lateral offset of the TMD dimer relative to the LBDs (**Fig. S10G**). Both classes show a TM6-TM6 inter-TMD interface with small differences in inter-TMD orientation (**Fig. S10H**), consistent with prior agonist- and PAM-bound full-length mGluR structures^37, 46–48, 57, 95–97^ (**Fig. S10H, I**). We observed density that is consistent with VU600 at both sides of the interface, suggesting a 2:2 binding stoichiometry (**Fig. S10J**,**K**). This PAM binding site is similar to that seen for a different PAM (VU036) in a prior mGluR4/G protein structure, although in that model only a single PAM per dimer is observed^48^ (**Fig. S10L**).

We next sought to structurally characterize mGluR8/β-arr1βCTD complexes using cryo-EM. While we observed multiple orientations and stoichiometries of β-arr1 in our negative stain analyses, cryo-EM of GPCR-β-arr1 complexes has been shown to only resolve core-bound β-arr1^21, 98, 99^. Indeed, extensive image processing resulted in a single class of particles (**Fig. 4F, S11A-D**) containing an active-like mGluR8 dimer and a density on the intracellular face with similar size, shape, and orientation to the 2:1 core-bound density from our negative stain EM experiments (**Fig. S11E**). This asymmetric arrestin-shaped density allows for unambiguous placement of a β-arr1 molecule with canonical interactions between the finger loop motif of β-arr1 and a specific TMD of the mGluR8 dimer (termed “chain A”) (**Fig.4F; Fig. S11F**). Additionally, this density is compatible with an AlphaFold2 prediction of the mGluR8/β-arr1 complex (**Fig. S11G**). While the density attributed to β-arr1 in our cryo-EM data was not at a resolution where features of the arrestin protomer aside from overall orientation were able to be discerned, the resolution of mGluR8 was similar to our “preactive” structures and allowed for changes in receptor conformation upon β-arr1 binding to be characterized (see below).

### Conformational changes of mGluR8 associated with transducer coupling

To enable a structural comparison of mGluR8 coupled to β-arr1 versus G proteins, we also determined a structure of L-AP-4 and VU600-bound mGluR8 in complex with Gα_i1_/β_1_/ψ_2_ heterotrimers (see Methods) (**Fig. 4G; Fig. S12**). mGluR8/Gα_i1_ showed an expected 2:1 receptor: heterotrimer stoichiometry and a similar overall orientation to prior G protein-bound mGluR structures (**Fig. S13A**).

Together, our pre-active, G protein-bound, and β-arr1-bound structures of mGluR8 allow for a comparison of structural changes associated with activation and desensitization. Like our pre-active structures, we observe active-like orientations of closed mGluR8 LBDs and compacted CRDs when bound to G protein or β-arr1 (**Fig. S13B, C**). We focused our analysis on changes at the level of the TMDs, as the structural changes induced by transducer coupling are most prominent near the interaction sites. Both β-arr1- and G protein-bound mGluR8 structures exhibit TM6-containing inter-TMD interfaces (**Fig. 4H**), consistent with both transducers stabilizing active states. Alignments of TMD dimers revealed more similar orientations for pre-active class 1 and β-arr1 versus pre-active class 2 and G protein (**Fig. S13D**). This may indicate that the two classes of pre-active structures represent sub-states that are biased toward binding a specific transducer protein. Subtle differences in the TMD dimer orientation are seen between β-arr1 and G protein structures, with a repositioning of chain A TM6 and a rotation of chain B relative to chain A (**Fig. 4H, I**). Inter-TMD distance measurements reveal asymmetric differences between structures (**Fig. S13E-G**), highlighting a modest G protein-induced tightening of the bottom chain A TM6-chain B TM7 interface and a modest β-arr1-induced relaxing of the same interface. Compared to pre-active structures where two PAM molecules are seen, the re-shaping of the TMD interface when bound to either G protein or β-arr1 result in the resolution of only one PAM molecule in each structure (**Fig. S13H**). Interestingly, the tighter inter-TMD interface seen for mGluR8/G protein results in PAM being located near TM1 of chain A while the PAM is seen on the opposite side of the dimer near TM1 of chain B in the mGluR8/β-arr1 structure.

In addition to changes in inter-subunit orientation, clear intra-subunit conformational differences were also observed in the TMD when bound to transducers. The most pronounced changes arise in chain A, the subunit which makes direct interactions with both β-arr1 and G proteins (**Fig. 4J; Fig. S13I)**. This includes a repositioning and lengthening of mGluR8 TM3, which is more pronounced in the presence of G protein compared to β-arr1 (**Fig. 4J**). This change results in a different orientation of ICL2 in the G protein and β-arr1 bound structures. In addition to changes to TM3, we observe an upward movement of TM6 in the mGluR8/G protein complex, reminiscent of what has been seen in an mGluR2/G protein structure^46^. Finally, we also observe a lengthening and different repositioning on the intracellular face of TM6 and TM7 between G protein and β-arr1 structures (**Fig. 4J**). Notably, these conformational changes do not include a canonical outward motion of TM6 as seen for activated family A and B GPCRs^100^ but has not been seen in family C GPCRs.

Chain B also showed subtle conformational changes in the presence of transducers, with the most drastic change occurring at TM3 where we observe similar lengthening in the presence of both G protein and β-arr1 (**Fig. S13J**). This corresponds to a more similar ICL2 orientation in both structures compared to what is seen in chain A. Conformational changes in chain B of the mGluR8/β-arr1 structure may hint at non-canonical interactions between the β-arr1 C-domain and mGluR8 that are not present in the more canonical finger-loop based interactions with mGluR8 chain A. These changes also suggest that a primary interaction with one TMD can lead to compensatory or modulatory conformational changes in the other subunit.

### Structural and computational analysis of mGluR8/β-arr1 complex dynamics

The positioning of β-arr1 relative to the homologous seven-helix TMD of various GPCRs shows a range of orientations across reported structures with mGluR8 displaying a unique x-y orientation in the plane parallel to the TMDs that is 90-150° shifted from that of the CB1R, β1AR, 5-HT2BR, and V2R and ∼180° opposite of that seen for the NTS1R (**Fig. 5A, B**). Interestingly, this x-y orientation is closest to that recently seen in mGluR3/β-arr1 structures^99^, suggesting a potential common arrangement for the mGluR subfamily. However, mGluR3 and mGluR8 still show a substantial >60° offset. Variability has also been observed in the orientation of β-arr1 relative to the plane of the membrane with some receptors (e.g. NTS1R, V2R) showing a sharp tilt z angle of <70° that may enable interaction between the β-arr1 C-edge and the membrane (**Fig. S13K**). In contrast, in our mGluR8 structural model, the β-arr1 C-edge is partially blocked from accessing the membrane by the other mGluR8 TMD and results in β-arr1 having a z angle of 74.6° (**Fig. 5A; Fig. S13K**).

**Fig. 5.**
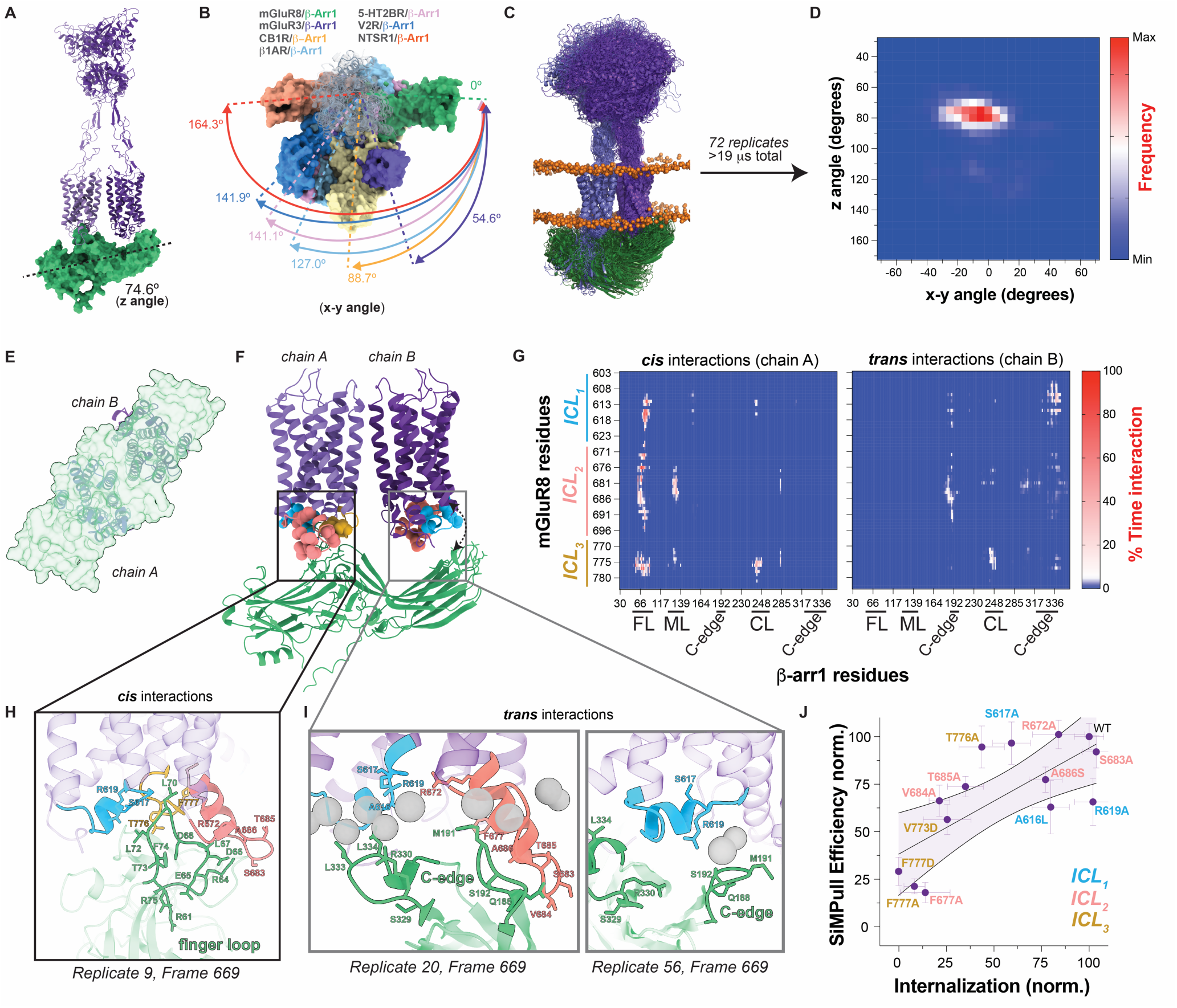
Structural, computational, and functional analysis of mGluR8/β-arr1 core coupling. **(A)** Model of mGluR8/β-arr1 bound to β-arr1 guided by cryo-EM density and AlphaFold2. The β-arr1 tilt z-angle is shown. **(B)** Overlay of the mGluR8/β-arr1 model with β-arr1 bound structures of family A GPCRs and mGluR3 (PDB: 8WU1, CB1R-β-arr1; 6TKO, β_1_AR-β-arr1; 7SRS, 5-HT2BR-β-arr1; 7R0C, V2R-β-arr1; 6UP7, NTSR1-β-arr1; 9II2, mGluR3-β-arr1, 2:2 stoichiometry). The different β-arr1 orientations are quantified by the x-y angle of rotation relative to that in mGluR8. **(C)** Snapshot showing overlay of the final mGluR8/β-arr1 conformation from 72 MD trajectories. **(D)** Heat map of x-y and z-angle of β-arr1 relative to mGluR8 across all MD simulations. **(E)** Cytoplasmic view of mGluR8/β-arr1 model showing that anchoring via core interactions with one subunit (light purple, left) enables a β-arr1 (translucent green) orientation that sterically occludes access to both subunits. **(F)** Side views showing mGluR8/β-arr1 interface from starting frame. Intracellular loop residues are shown as spheres in blue (ICL1), pink (ICL2), and gold (ICL3). **(G)** Heat map summarizing interactions between chain A (left) and chain B (right) of mGluR8 across all MD simulations. **(H)** Snapshot from MD simulations highlighting key residues involved in *cis* interactions between the β-arr1 finger loop and mGluR8 chain A. **(I)** Snapshots from MD simulations highlighting two different interaction modes between the β-arr1 C-edge loops and the TMD of mGluR8 chain B and membrane lipids (grey spheres). **(J)** Scatter plot showing correlation between effects of mutations on β-arr1-βCTD co-localization in SiMPull (y-axis) and internalization in cell-based surface labeling assay (x-axis). A linear fit is shown with the 95% confidence interval shaded. All data shown as mean ± SEM.

To better understand the molecular details of the mGluR8/β-arr1 interaction, we performed molecular dynamics simulations using a cryo-EM based structural model of the mGluR8/β-arr1 complex (see Methods). In total, we ran 72 independent replicas of the mGluR8/β-arr1 complex embedded in a model plasma membrane, resulting in over 19 µs of total simulation time. Across all simulations the mGluR8 domains remain in an active-like conformation (**Fig. 5C; Fig. S14A, B**) containing closed-closed/active LBD dimers (**Fig. S14C**) and a TM6-TM6 inter-TMD interface (**Fig. S14A, C**). However, we observed a tilting motion of the LBD relative to the TMD (**Fig. 5C; Fig. S14A**), revealing large-scale conformational flexibility of mGluR ECDs. This domain flexibility is consistent with the variable LBD tilt seen in our cryo-EM structures of mGluR8 (**Fig. 4E-G**).

β-arr1 remained stably bound via its finger loop motif to the TMD core of mGluR8 in a range of active-like conformations (**Fig. S15A, B**) throughout the simulations but showed positional dynamics in both X-Y and Z planes relative to the mGluR8 TMDs (**Fig. 5D**; **Fig. S15C-H**). This conformational heterogeneity is consistent with the limited resolution of the β-arr1 density in our cryo-EM maps. Importantly, the β-arr1 orientation resolved in our cryo-EM model, in terms of the x-y and z-angle falls in the middle of the dominant orientation of the β-arr1 in the complex (**Fig. 5D**). We also observed a subset of simulations (13/72) where the β-arr1 C-edge eventually falls away from the TMD, resulting in a subpopulation of simulation frames where the z-angle is >100° and further highlighting the dynamics of the system (**Fig. S15F-H**).

Our cryo-EM structure and MD simulations of a 2:1 core-bound mGluR8/β-arr1 reveal a primary interaction where the orientation of β-arr1 relative to mGluR8 is such that the cytoplasmic surface of both TMDs is largely occluded (**Fig. 5E**). This suggests a simple steric mechanism for β-arr-mediated desensitization of mGluR8 that would provide an efficient mechanism for preventing G protein binding to the receptor dimer, effectively preventing canonical signal initiation. In addition, this orientation would preclude the binding of a second β-arr molecule to the TMD dimer core without either a large rearrangement of the β-arr or a major inter-subunit rotation of the TMD. The core binding mode of β-arr1 observed in our structure does not preclude a second β-arr1 molecule binding to the same receptor dimer via the second mGluR8 CTD in a tail-only mode. This configuration may be compatible with our negative stain data of 2:2 complexes (**Fig. 4D**). Interestingly, 2:1 and 2:2 mGluR3/β-arr1 structures were recently reported, with both β-arr1 molecules in a core-bound mode in the 2:2 model^99^. In contrast to the active-like TM6 interface observed here for mGluR8, in this structure an inactive-like TM3-containing inter-TMD interface was observed, providing sufficient space to avoid a steric clash between β-arr1 protomers.

To better understand the residues mediating the mGluR8/β-arr1 interaction, a closer analysis of interactions between β-arr1 and the mGluR8 TMD in our model and simulations was performed. A contact heat map was generated for interactions between β-arr1 and both mGluR8 TMDs which revealed interactions between the β-arr1 finger loop and residues in ICL1, ICL2, and ICL3 of the receptor chain A (**Fig. 5G**). The finger loop binding site is comprised primarily of intracellular loops and is highly overlapping with what is seen for the α5 helix of Gα_i1_ in our G protein-bound mGluR8 structure (**Fig. 5G, H; Fig. S16A, B**). Consistent with coupling via these interactions, we observed reduced structural variability in this TMD subunit (chain A) in our simulations (**Fig. S14B**) and a consistent but dynamic interface between the finger loop and mGluR8 (**Fig. S16C, D, F**). We also observed interactions between the β-arr1 middle and C-loops and mGluR8 chain A ICL 2 and 3, respectively (**Fig. S16E-G**), which may contribute to the stability of the core mGluR8/β-arr1 interaction. Finally, in trajectories with large β-arr1 z-angles (> 90°), we observed that interactions between residues in the β-arr1 N-lobe and the membrane may stabilize this state in a lipid-dependent manner (**Fig S16H, I**). These interactions may be disfavored in our micelle-solubilized, purified mGluR8/β-arr1 complexes which may explain why we do not identify this β-arr1 orientation in our EM experiments.

Owing to the orientation of β-arr1 relative to the mGluR8 dimer, the β-arr1 C-edge comes into proximity with both ICL1 and ICL2, and to a lesser extent ICL3, of the mGluR8 chain B TMD (**Fig. 5F, G, I, Fig S16J, K)**. We also observe extensive interactions between β-arr1and the plasma membrane in many of our replicas (**Fig. S16L-N**). ICL1/C-edge interactions are mediated electrostatically and often recruit a negatively charged lipid headgroup (**Fig. 5I**, left) while ICL2 interactions are formed via a long, hydrophobic interface (**Fig. 5I**, right). We also observe C-edge loops embedding into the membrane in some replicas without apparent interactions with mGluR8 (**Fig. S16K, L**) and with specific lipid preferences (**Fig. S16N**). This observation is in line with recent biophysical studies showing β-arr C-edge/membrane interactions^29, 101^and further highlights the apparent configurational diversity exhibited by β-arr1 while maintaining the canonical finger loop-based interaction with mGluR8.

To experimentally probe the mGluR8 TMD/β-arr1 core interaction observed in our model and simulations, we individually mutated putative interfacial residues in ICL1, ICL2, and ICL3. Using SiMPull, we found impaired β-arr1-βCTD-Halo colocalization with mGluR8 mutants across all loops with the largest effects for F677A and F777A in ICL2 and ICL3, respectively (**Fig. 5J; Fig. S17A**). Interestingly, the melanoma-associated mutation V773D in ICL3^102^ also substantially impaired interaction with β-arr1 (**Fig. 5J; Fig. S17A**). Similar effects of the mutations were observed using our surface labeling internalization assay (**Fig. 5J; Fig. S17B, C**) with a clear correlation between the two assays (**Fig. 5J**). Results were most variable and subtle for mutations in ICL1, suggesting that these interactions are secondary in importance compared to the primary interactions with ICL2 and ICL3, in line with what has been previously observed in mutational analysis of mGluR/G protein coupling^47^.

### β-arr1 can couple to mGluR8 in both *cis* and *trans* configurations

The involvement of both the mGluR8 tail and core in β-arr1 coupling, and the orientation of β-arr1 found in our structural studies, raises questions about the possible configurations of mGluR8/β-arr1 complexes. Specifically, can coupling occur in via interactions of β-arr1 with the tail and TMD core of the same mGluR8 subunit (termed *cis*), and/or via interactions of β-arr1 with the tail of one subunit and the TMD core of the other (termed *trans*)? Notably, the active-like, β-arr1-bound mGluR8 structure brings the beginning of the CTD to a central site within the inter-TM6 interface. Also, the mGluR8 S/T-rich region is located at the end of the CTD, which could allow for the tail-bound arrestin-mGluR8 complex to maintain a high degree of conformational flexibility. Together, the structural features of arrestin-bound mGluR8 position it such that it is conceivable that arrestin coupling to the TMD core of either subunit can occur while bound to the CTD.

To test this, we took advantage of mutations described above that strongly impair either tail interaction (“11xA”) or core interaction (F777D) to prevent mGluR8/β-arr1 co-internalization (**Fig. 5A; Fig. S18A**). We first co-expressed SNAP-tagged mGluR8-F777D with untagged mGluR8-11xA. These expression conditions should produce a mix of SNAP-tagged F777D homodimers and F777D/11xA heterodimers, as well as 11xA homodimers which are unlabeled. Given that 11xA homodimers show no internalization (**Fig. 2B**), we reasoned that any visible intracellular fluorescence or drop in surface fluorescence in the surface labeling assay would indicate heterodimer internalization. Indeed, we observed clear internalization both in cell imaging and surface labeling internalization assays for this mix of receptor subunits (**Fig. 5A; Fig. S18A**). This finding supports the ability of *trans* interactions whereby β-arr binds to the CTD of the F777D subunit and the core of the 11xA subunit to mediate β-arr-mediated internalization. Similar results were observed upon co-expression of SNAP-mGluR8-11xA with untagged mGluR8-F777D (**Fig. 5A; Fig. S18A**). Supporting *cis* interactions of β-arr1 with the CTD and core of the same subunit, co-expression with wild type untagged mGluR8 enabled internalization of SNAP-mGluR8-F777D-11xA (**Fig. 5A; Fig. S18A**).

To directly detect and compare the propensities of *cis* and *trans* mGluR8/β-arr1 interactions, we turned to SiMPull. First, we generated an HA-tagged mGluR8-11xA construct that did not contain a SNAP-tag and should be unable to pulldown any β-arr1-Halo based on our prior results (**Fig. 2A**). To detect *trans* interactions, we tested the ability of this construct to pulldown with WT β-arr1 upon co-expression with SNAP-mGluR8-F777D and observed substantial ∼20% colocalization (**Fig. 6B**), presumably via tail interactions with the F777D subunit. Importantly, the colocalization percentage for this *trans*-coupling mGluR8 heterodimer was larger than pulldown for homodimeric mGluR8-F777D (**Fig. 3C**). To detect *cis* interactions, we produced an HA-mGluR8-F777D-11xA construct and tested its ability to pulldown with β-arr1 upon co-expression with wild type SNAP-mGluR8, and, again, observed substantial colocalization (**Fig. 6C**), suggesting a complex where both tail and core interactions occur via the WT subunit. Importantly, in these experiments we are only able to detect heterodimers (F777D/11xA or F777D-11xA/WT) since only one subunit is labeled with HA for pulldown and the other subunit contains a SNAP-tag for fluorophore conjugation. Quantification of *trans* and *cis* SiMPull conditions showed clear colocalization in both cases with enhanced efficiency in *cis* (**Fig. 6D**), suggesting a weak preference for this mode of tail and core interactions.

**Fig. 6.**
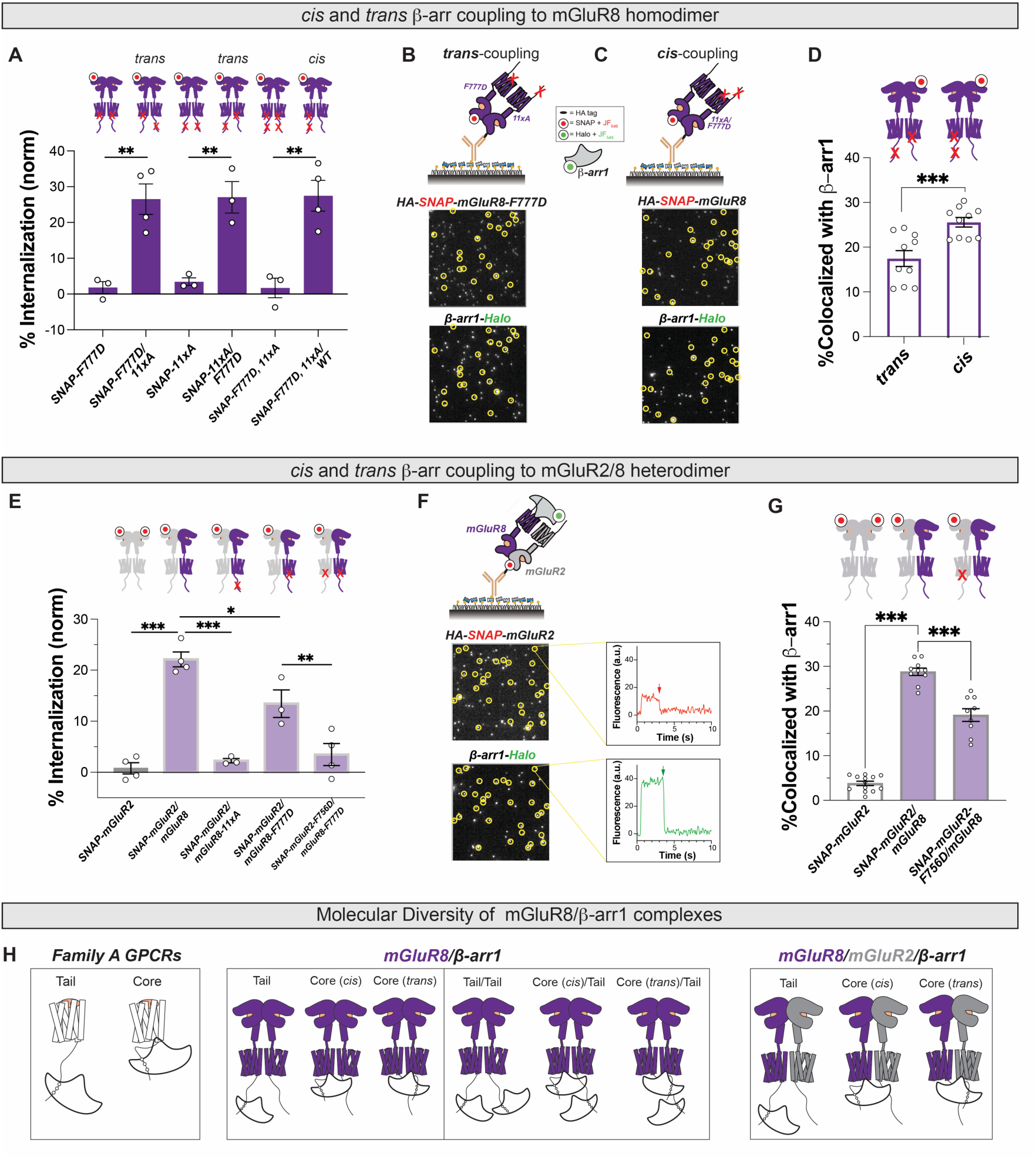
β-arr coupling in *cis* and *trans* configurations to homo and heterodimeric mGluR8. **(A)** Surface labeling assay internalization data showing evidence for *trans* (SNAP-F777D/11xA; SNAP-11xA/F777D) and *cis* (SNAP-F777D,11xA/WT) β-arr1 coupling driving mGluR8 internalization. **(B-D)** Representative SiMPull images and summary quantification showing evidence for *cis* and *trans* mGluR8/β-arr1 coupling with higher efficiency for *cis* compared to *trans* **(D)**. **(E)** Surface labeling assay internalization data showing evidence for mGluR2/8 heterodimer internalization via mGluR8 CTD coupling and core coupling to either subunit. **(F)** Representative SiMPull image (left) and photobleaching step traces (right) showing the formation of 1:1:1 mGluR2: mGluR8: β -arr1 complexes. **(G)** Quantification of SiMPull showing evidence for mGluR2/8/ β -arr1 complexes that are sensitive to core mutation (F756D) in mGluR2. **(G)** Schematic of proposed modes of β-arr1 coupling for family A GPCRs (2 modes), mGluR8 (6 modes), and mGluR2/8 (3 modes) highlighting the diversity of potential conformations of dimeric GPCR/β-arr complexes that can exist. 1-way ANOVA is used in **(A, E, G)** and unpaired t-test is used in **(D**). *, p<0.05; **, p<0.01; ***, p<0.001. All data shown as mean ± SEM.

The ability of β-arr1 to couple to mGluRs in either *cis* or *trans* configurations may be particularly relevant for heterodimeric mGluRs which introduce asymmetry into the cytoplasmic face of the receptor and may be regulated in complex ways by subunit-specific pharmacology or post-translational modifications. We focused on mGluR2/8 heterodimers as these combine a strong β-arr coupled mGluR (mGluR8) with a β-arr resistant mGluR (mGluR2) and we have recently shown that these heterodimers undergo β-arr dependent endocytosis and lysosomal degradation^63^. We first confirmed that co-expression with untagged mGluR8 enabled internalization of SNAP-mGluR2 (**Fig. 6E; Fig. S18B**). Introducing the 11xA mutation into mGluR8 abolished mGluR2 internalization but introducing the F777D mutation into mGluR8 only partially impaired mGluR2 internalization (**Fig. 6E; Fig. S18B**), suggesting that while mGluR8 tail coupling is strictly required, mGluR8 core coupling is not. Internalization was fully abolished when the equivalent mutation, F756D, was also introduced into mGluR2 (**Fig. 6E; Fig. S18B**), suggesting that both *cis* and *trans* mGluR2/8 β-arr coupling can occur via the mGluR8 tail and either the mGluR2 or mGluR8 core. Using SiMPull we found that co-expression of SNAP-mGluR8 and HA-SNAP-mGluR2 enables colocalization with β-arr1-Halo (**Fig. 6F, G**). Importantly, the vast majority of colocalized β-arr1-Halo showed single-step photobleaching (**Fig. 6F, S18C**), consistent with a 1:1:1 mGluR2: mGluR8: β-arr1 complex. Finally, we observed a decrease in β-arr1-Halo colocalization when we introduced the F756D mutation into mGluR2, further supporting both *cis* and *trans* β-arr1 coupling to mGluR2/8 heterodimers (**Fig. 6G**).

Together with our EM data showing core and tail-bound mGluR8/β-arr1 coupling (**Fig. 4**) and the steric constraints of the core-bound mGluR8 structure (**Fig. 5E**), these data allow us to propose 6 distinct modes of mGluR/β-arr1 coupling with either 2:1 or 2:2 receptor/arrestin stoichiometries which contrast with the limited repertoire of 1:1 family A GPCR/b-arr1 coupling (**Fig. 6H**).

### mGluR8 can simultaneously bind β-arr1 and β-arr2 to form megacomplexes

While β-arr1 has served as the model β-arr subtype for many structural and biophysical studies, β-arr2 also strongly contributes to GPCR regulation. As β-arr2 has known structural and functional differences compared to β-arr1^24, 33–36^, we first confirmed that β-arr2-Halo co-internalizes with both mGluR8 and V2R upon agonist stimulation (**Fig. S19A, B**). Using SiMPull, we observed clear pulldown of WT β-arr2 by mGluR8 and V2R, but not mGluR2, although the efficiency was reduced in comparison to WT β-arr1 (**Fig. 7A-C**). mGluR8-mediated pulldown of WT β-arr2 was dramatically reduced with the 11xA or F777D mutations (**Fig. S19C**), suggesting contributions from both tail and core interactions. To determine whether we could stabilize the pulldown of β-arr2, we produced two “pre-activated” β-arr2-Halo constructs^36^: one containing 3 alanine mutations (I386A, V387A, F388A) in the proximal CTD of β-arr2 (β-arr2-3A) and one containing these mutations and a subsequent truncation of the β-arr2 CTD (β-arr2-3A-ΔCTD). Both β-arr2 variants had little impact on colocalization percentage for mGluR8 but increased pulldown for V2R (**Fig. 7C**). The differential effect of pre-activating β-arr2 modification across receptor subtypes highlights the complexity and receptor subtype-specificity of GPCR/β-arr coupling. Photobleaching step analysis showed both 2:1 and 2:2 mGluR8/β-arr2 complexes (**Fig. 7B, D**) indicating that the same stoichiometries are observed for complexes of mGluR8 with β-arr1 or β-arr2. As was observed with β-arr1, V2R/β-arr2 complexes showed a 1:1 stoichiometry (**Fig. 7D**). We further confirmed the potential for 2:2 mGluR8/β-arr2 assembly by observing substantial colocalization between β-arr2-Halo molecules when labeled with two different colors upon pulldown by HA-mGluR8 (**Fig. S19D**). Finally, we purified mGluR8/β-arr2-3A-ΔCTD and performed negative stain EM to assess whether mGluR8/β-arr2 complexes exhibit similar structural orientations as mGluR8/β-arr1ΔCTD complexes including both core- and tail-bound orientations of β-arr2 and both 2:1 and 2:1 mGluR8:β-arr2 stoichiometries in purified complexes (**Fig. 7E, F; Fig. S20**). Together, we find stoichiometric and conformational similarities between mGluR8 complexes with both β-arr1 and β-arr2, while also exhibiting apparent β-arr isoform specific differences in stability of the complexes.

**Fig. 7.**
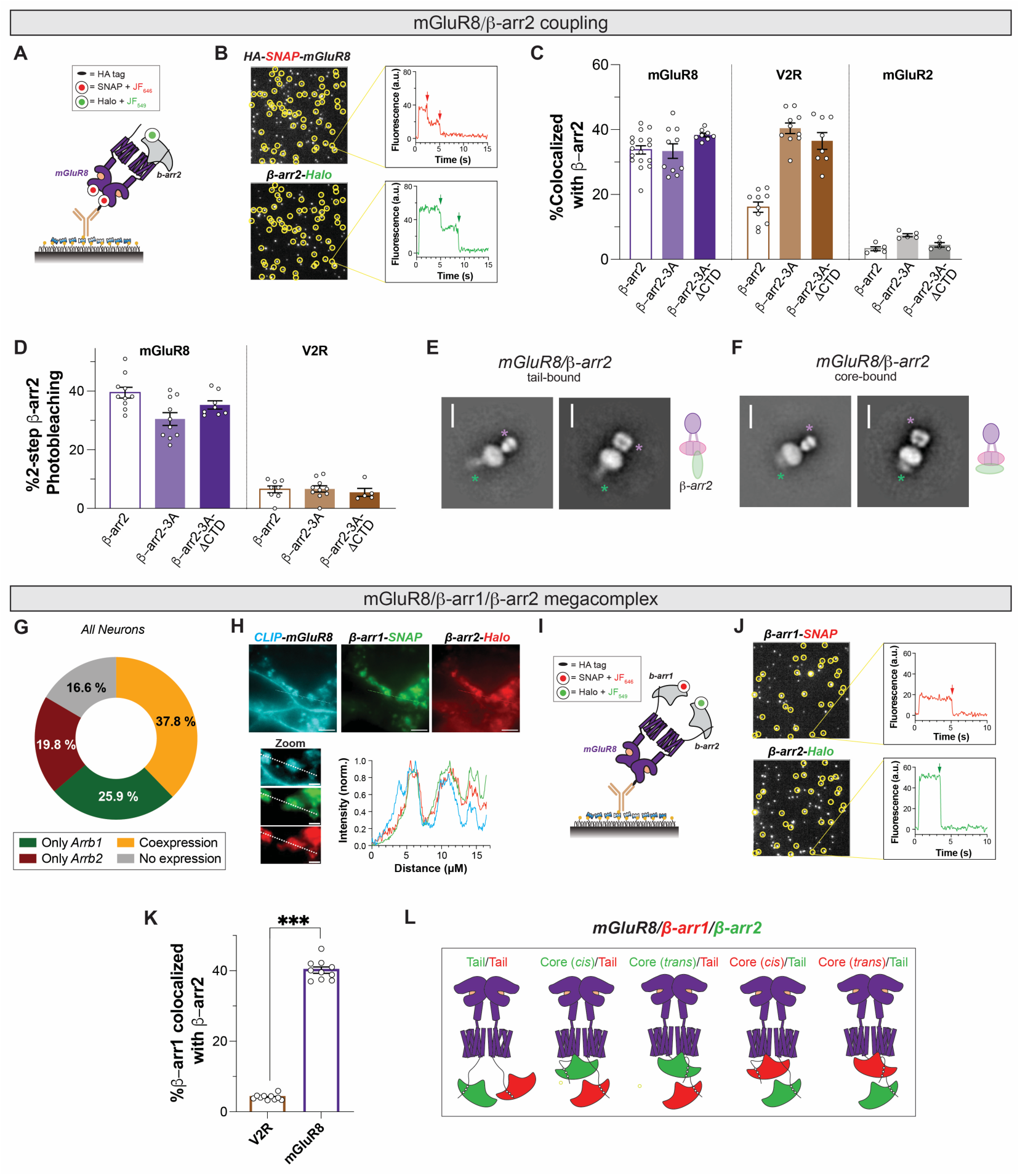
Single molecule analysis of assembly with β-arr2 and formation of mGluR8/ β-arr1/ β-arr2 megacomplexes. **(A-B)** Schematic (A) and representative image (B) showing colocalization and representative bleaching steps of mGluR8 and β-arr2. **(C)** Colocalization percentages of β-arr2 and β-arr2 variants across a panel of GPCRs. **(D)** Summary graph showing substantial formation of complexes with 2 β-arr2 bleaching steps for mGluR8, but not V2R. **(E-F)** Negative stain EM 2-D classes showing tail- (E) and core- (F) bound forms of mGluR8/β-arr2. **(G)** Pie chart showing proportion of neurons expressing 0, 1, or both β-arr genes determined from single cell RNA sequencing data using a cutoff of 5 counts per million. **(H)** Representative image and line scan showing colocalization of intracellular mGluR8, β-arr1, and β-arr2. **(I-J)** Schematic (I) and representative image (J) showing co-localization of β-arr1 and β-arr2 upon pulldown via mGluR8. **(K)** Summary graph showing β-arr1/β-arr2 colocalization upon pulldown by mGluR8, but not V2R. **(L)** Schematic of proposed modes of simultaneous β-arr1 and β-arr2 coupling to mGluR8. Points represent individual movies (C, D, K). GRK2-CaaX was co-expressed for all SiMPull experiments shown in this figure. Unpaired t-test is used in (K). ***, p<0.001. All data shown as mean ± SEM.

As mGluR8 interacts with both β-arr1 and β-arr2 and can form a 2:2 stoichiometry, we next asked if a single mGluR8 dimer can simultaneously bind both β-arr1 and β-arr2. We first analyzed single cell RNA sequencing (scRNAseq) data from the mouse anterior cingulate cortex^103^ to evaluate whether β-arr1 (gene name: *Arrb1*) and β-arr2 (gene name: *Arrb2*) are co-expressed in the same neurons. We found subtle differences in β-arr1 versus β-arr2 expression across neuronal subtypes (**Fig. S19E**) with a consistent trend towards more β-arr1 expression in most classes. Overall, 37.8% of neurons showed co-expression of β-arr1 and β-arr2 (**Fig. 7G**). Notably, a wide range of RNA ratios between mGluR8, β-arr1, and β-arr2 is seen across neurons with a large proportion (∼24% of the total and ∼40% of mGluR8 expressing neurons) expressing all three (**Fig. S19F, G**), raising the possibility that under biologically relevant conditions β-arr1 and β-arr2 can either compete for the same GPCR or bind in a combinatorial fashion. Accordingly, live cell imaging revealed endosomal co-localization of CLIP-mGluR8, β-arr1-SNAP, and β-arr2-Halo (**Fig. 7H**), pointing towards this combinatorial possibility. To address this directly, we performed SiMPull from cells expressing HA-mGluR8 lacking a SNAP tag along with GRK2-CaaX (**Fig. 7I**) and found a large proportion of spots with colocalized β-arr1-SNAP and β-arr2-Halo (**Fig. 7J, K**), confirming the presence of an mGluR8/β-arr1/β-arr2 megacomplex. Importantly, β-arr1-SNAP and β-arr2-Halo channels showed a single bleaching step in each channel, consistent with a 2:1:1 complex (**Fig. 7J**). In contrast, negligible background levels of β-arr1-SNAP and β-arr2-Halo colocalization were observed with pulldown of V2R (**Fig. 7K; Fig. S19H**). This result represents, to our knowledge, the first example of two different arrestin subtypes simultaneously interacting with a single GPCR and enables the proposal of a wide range of β-arr2 containing mGluR complexes (**Fig. 7L**), further expanding the molecular diversity of mGluR/arrestin coupling.

## Discussion

Our integrative biophysical approach reveals major complexity in mGluR/β-arr coupling, suggesting a high degree of tuneability and flexibility to match the myriad roles of mGluRs in regulating synaptic transmission and plasticity^71, 72, 104^. The ability of mGluRs to couple to either one or two copies of either β-arr1 or β-arr2 in tail or *cis* and *trans* core-binding modes may serve to impart different extents or speeds of desensitization, endocytosis, intracellular trafficking, and/or arrestin-dependent signaling. Combined with the ability of mGluR heterodimers to couple to β-arrs in multiple configurations and the possibilities of mGluR/β-arr1/β-arr2 megacomplex, we propose at least 20 distinct modes of mGluR8/β-arr coupling, contrasting with the limited repertoire of 1:1 family A GPCR/ β-arr coupling. It is likely that the specific mode of β-arr binding that predominates in each setting depends on the degree or timing of mGluR activation by synaptic glutamate or exogenous ligands and the given cellular context in terms of the β-arr1 and β-arr2 expression levels, subcellular localization, cell morphology, and the presence of other regulatory proteins (e.g. GRKs).

Our study supports an increasing appreciation of the functional specialization of mGluR subtypes. Our data correlate well with prior studies that found that mGluR2 is resistant, while mGluR3 undergoes transient β-arr coupling on the cell surface followed by recycling, and mGluR8 undergoes co-internalization followed by lysosomal degradation^62, 63^. Our observations that mGluR8 has a steeper dependence on GRK expression but shows a higher complex stability than mGluR3 (**Fig. 1**), suggest that mGluR3 is specialized for rapid, reversible desensitization while mGluR8 is optimized for longer-lasting downregulation after elevated or extended degrees of activation, potentially allowing each receptor to regulate distinct forms of synaptic plasticity and metaplasticity. Interestingly, mGluR5 showed complex formation with β-arr1- βCTD in SiMPull despite evidence that it undergoes β-arr-independent endocytosis^62, 105–107^. This suggests that relatively weak β-arr coupling may play non-trafficking roles in tuning or initiating mGluR5 signaling, in line with studies showing an impairment of mGluR5-mediated long-term depression following β-arr2 knockout^108^. In contrast, mGluR7 showed weak complex formation with β-arr1 in SiMPull despite evidence that it undergoes β-arr-dependent internalization and degradation in neurons^64^. This prior work revealed a strong dependence on ubiquitination of mGluR7 for its degradation and, together with our work, suggests that β-arr-dependent GPCR degradation may not strictly require the formation of a tight complex with β-arrs. The distinct modes of β-arr coupling in terms of complex stability and intracellular trafficking uncovered here suggest distinct molecular and/or structural mechanisms across subtypes. Notably, our biophysical and structural findings for mGluR8/β-arr complexes differ from a recent study on mGluR3/β-arr complexes which found that β-arr1 stabilizes an inactive-like conformation of mGluR3 and that two β-arr1 molecules can simultaneously bind to the core of adjacent TMDs^99^, supporting the idea that different mGluRs utilize different modes of β-arr interactions.

Structural analysis of mGluR8 via cryo EM and MD simulations allowed us to visualize and model the orientation of β-arr1 on agonist and PAM-bound, full-length, human, wild-type mGluR8, providing a glimpse into a family C GPCR/β-arr complex. The unique orientation of β-arr1 relative to the TMD of mGluR8 is not well-aligned with existing GPCR/β-arr1 structures, extending our appreciation of the flexibility/dynamics involved in β-arr coupling to GPCRs. Intriguingly, the single core-bound β-arr1 seen with mGluR8 is well-positioned to both occlude G protein binding to the subunit that it directly interacts and to the adjacent subunit via its C-lobe. This bridging of two GPCR TMD hints at the ability of a single β-arr to desensitize two monomeric GPCRs by a similar mechanism which has been proposed for rhodopsin^109^ and the mu-opioid receptor^110^. The active conformations observed for mGluR8 in our complex structure, with repositioning of intracellular loops and the maintenance of a TM6-containing interface, supports our smFRET analysis which revealed mGluR8 active-state stabilization by β-arr1. These findings suggest that, analogously to family A GPCRs that have been shown to be stabilized in active states by β-arrs^29, 111, 112^, both G proteins and β-arrs can couple to and stabilize similar active conformations in mGluRs. Subtle differences in both TMD conformation and TMD dimer orientation between β-arr1 and G protein-bound mGluR8 structures suggest unique active states and, thus, the possibility of developing biased ligands that preferentially promote or inhibit arrestin versus G protein coupling.

The molecular diversity of mGluR/β-arr coupling modes uncovered in this study motivates further structural analyses across mGluR and β-arr subtypes. Recent studies have shown that β-arr2 can adopt distinct family A GPCR binding modes^24, 36^, raising the question of how it couples to mGluRs and how mGluR/β-arr1/β-arr2 megacomplexes may be assembled. Notably, a seemingly similar mGluR8 core-bound orientation was seen for β-arr2 compared to β-arr1 in our negative stain EM data, buthigher resolution structural data is needed for a more detailed comparison of mGluR8 interaction with β-arr1 versus β-arr2. Intriguingly, our findings raise the possibility of megacomplexes between an mGluR, β-arrs, and G proteins, as variations of this have been seen in a family A GPCRs and may contribute to endosomal signaling^113–116^. Notably, the core-bound model of mGluR8/β-arr1 we observed here would be incompatible with simultaneous binding of β-arr1 and a G protein heterotrimer without major reconfiguration. However, G protein binding may be compatible with tail-bound β-arr or with alternative conformations, such as the high β-arr1 z-angle binding mode observed in our MD simulations. As we have demonstrated mGluR and GRK subtype dependence on β-arr coupling, further work is also required to understand the structural basis of GRK/mGluR interaction. Finally, the CTDs of mGluRs are known to interact with myriad other proteins at the synapse, including calmodulin, Munc18, PICK1, and Homer^71, 117–121^, raising the question of how such interactions may prevent or potentiate β-arr binding and further sculpt mGluR regulation.

## Methods

### Molecular Biology

All SNAP- or CLIP-tagged mGluR2 (rat), mGluR3 (rat), mGluR5 (human), mGluR7 (rat), mGluR8 (human), V2R (human), β2AR (human) were generated with an N-terminal signal peptide from rat mGluR5 followed by variable combinations of HA-, SNAP-, FLAG-, or CLIP in the pRK5 vector as previously described^62^. Untagged bovine GRK2 (#14691), GRK2-CaaX (#166224), and GRK5 (#14690), were purchased from Addgene. Point mutations were introduced using PCR-based site-directed mutagenesis. Rat β-Arr1–Halo, β-Arr1–SNAP, and β-Arr2–Halo were generated by replacing YFP from β-arr1–YFP (Addgene) or β-arr2–YFP (Addgene, #36917) with the Halo or SNAP tag using a Gibson Assembly Cloning Kit. β-arr1-ΔCTD-Halo constructs were created by removing residues 383-418 from the WT β-Arr1–Halo plasmid using In-Fusion cloning (Takara Bio). mGluR8-ΔCTD was created by introducing a STOP codon in position 845 by PCR-based site-directed mutagenesis. β-Arr2-3A constructs were created by mutating residues 386-388 to alanine and β-arr2-3A-ΔCTD constructs were made by introducing a STOP codon after residue 393 using In-Fusion cloning. For cryo-EM experiments, cDNA for human mGluR8 was cloned into the pEZT-BM BacMam expression vector ^122^ and the N-terminus was fused to an HA and FLAG (DYKDDDD) tag followed by a 3-alanine linker (AAA).

### Cell culture and transfection

HEK 293T (ATCC: CRL-3216) cells were grown and maintained in Dulbecco’s Modified Eagle’s Medium (Corning) supplemented with 10% fetal bovine serum (Thermo Fisher) and cultured at 37°C/5% CO_2_. For live cell imaging and single molecule imaging experiments, cells were seeded on poly-L-lysine (Sigma) coated 18 mm glass coverslips in 12-well plates and transfected using Lipofectamine 2000 (Thermo Fisher Scientific). For cell imaging experiments (e.g. Fig. 1A, B), cells were transfected with 700 ng each of receptor, β-arr, and GRK2 per well. SiMPull experiments also used 700 ng of each plasmid per well. For some SiMPull experiments, GRK2 was replaced with GRK2-CaaX or GRK5 or omitted. For surface labeling assay experiments (e.g. Fig. S5B), only receptor was transfected. For experiments with expression of two receptor constructs, a 1:1 ratio was maintained with 700 ng of each receptor. For β-arr co-expression experiments a 1:2 ratio of β-arr1: β-arr2 was used. Six hours after transfection with mGluRs, media was replaced and supplemented with the antagonist LY341495 until the time of the experiment (5 µM for mGluR2 and mGluR3; 50 µM for mGluR7 and mGluR8) to maintain cell health.

### Live cell fluorescence imaging

For live cell imaging of receptor internalization/β-arr localization, 24-48 hours post-transfection cells were labeled for 30 minutes at 37°C with 1 µM of the appropriate SNAP (SNAP-Surface546 from NEB for SNAP-tagged receptors or custom-made permeable SNAP-JF_549_ for β-arr1–SNAP), CLIP (CLIP-Surface488, from NEB), or Halo (permeable JF_646_ from Promega) dyes in extracellular buffer (EX) containing (in mM): 10 HEPES, 135 NaCl, 5.4 KCl, 2 CaCl₂, and 1 MgCl₂, pH 7.4. After labeling, cells were washed twice with fresh EX to remove excess fluorophore and incubated in EX containing different agonists (1 mM Glu for mGluR2 and mGluR8, 100 µM Glu for mGluR3, 10 mM Glu for mGluR7, 10 µM Vasopressin for V2R, 10 µM Isoproterenol for β2AR) for 30 min. Images were acquired using an Olympus IX83 inverted microscope equipped with scientific Complementary Metal-Oxide Semiconductor (sCMOS) ORCA Flash4v3.0 camera (Hamamatsu) and a 100X immersion oil objective (NA 1.45). Fluorophores were excited with 488 nm, 561 nm, and/or 640 nm lasers in widefield mode.

mGluR internalization was assessed using a surface labeling assay as previously described^37, 62, 63^. Briefly, 24 hours post-transfection of SNAP-mGluR constructs, cells were incubated for 60 min at 37°C in warm DMEM + 10% FBS with either antagonist (50 µM LY341495) or agonist (1 mM Glu) for each condition. Subsequently, cells were labeled with 1 µM SNAP Surface Alexa 546 dye (NEB) in EX for 20 min at RT. Images were captured using an Olympus inverted microscope with a 60X immersion oil objective (NA 1.49). At least 10 images per condition were obtained per day, and the experiment was repeated for a minimum of three independent biological replicates. Quantification of remaining surface receptors was performed using the Li algorithm threshold method in ImageJ (Fiji). Background fluorescence mean intensity was subtracted from the mean intensity of fluorescent cells, and these values were normalized to the antagonist condition per day and construct. Percent internalization was calculated by normalizing the image values of the agonist condition to the antagonist condition, transforming these values into percentages, and averaging among all images per day. Each mean value represented an independent data point.

mGluR degradation was assessed as previously described^63^. Similarly to the surface labeling assay, 24 hrs post-transfection cells were incubated for 60 min at 37°C in complete DMEM containing 50 µM LY341495 or 1 mM Glu for each mutant (and WT as control). In this case, cells were labelled in EX buffer with a cell permeable dye, BG-Janelia Fluor 549 (1 µM) for 30 min at 37°C, followed by a 30 min wash. Images were acquired and analyzed using the same protocol described above for the surface labelling assay.

### Single Molecule Pulldown (SiMPull)

SiMPull measurements were performed as previously described^63, 68^ with minor modifications. Forty hours post-transfection (see details above), HEK 293T cells were washed with EX buffer, and then labeled for 1 hour at 37° C with 1 μM CA-JF_549_ (permeable; for Halo; Promega) and 1 μM SBG-JF_646_ (impermeable; for SNAP; custom synthesized as previously^123^). For experiments with dual labeling of β-arr1-Halo (Fig. S2L, M) or β-arr2-Halo (Fig. S7C), cells were simultaneously labeled with 1 μM CA-JF_549_ and 1 μM CA-JF_646_ (permeable; for Halo; Promega). For experiments with dual labeling of β-arr1-SNAP and β-arr2-Halo (Fig. 7I-K), cells were simultaneously labeled with 1 μM BG-JF_646_ (permeable; for SNAP; custom synthesized as previously^123^) and 1 μM CA-JF_549._ Following labeling, cells were washed with EX buffer before the incubation of agonist (1 mM Glu for mGluR2, mGluR3, mGluR5, mGluR8; 10 mM Glu for mGluR7; 10 µM Vasopressin for V2R; 10 µM Isoproterenol for β2AR) for 45 minutes at 37° C. Cells were then gently harvested in phosphate-buffered saline (0 Ca^2+^, 0 Mg^2+^) and pelleted at 10,000 rpm at 4°C for 5 minutes. Receptors were isolated via gentle lysis using 0.5% Lauryl Maltose Neopentyl Glycol (LMNG) + 0.05% Cholesteryl hemisuccinate (CHS) (Anatrace) plus protease inhibitor cocktail (Thermo Fisher Scientific) in Lysis Buffer (10 mM Tris HCl, 150 mM NaCl, 1 mM EDTA, pH 8.0) at 4°C for 1 hour. After lysis, samples were centrifuged at 17,000*g* for 20 min at 4°C, and supernatants were collected. A microflow chamber was prepared using a glass coverslip and quartz slide passivated with mPEG-

SVA and biotinylated-PEG (molecular weight: 5000; 50:1 molar ratio; Laysan Bio). Before each experiment, each chamber was washed with T50 buffer (50 mM NaCl and 10 mM tris, pH 8.0), incubated with NeutrAvidin (0.2 mg/mL) in T50 buffer for 5 minutes, followed by incubation with biotinylated anti-HA antibody (0.002 mg/ml; ab26228, Abcam) in T50 buffer for 20 minutes. Flow chambers were washed with T50 buffer after incubations with both NeutrAvidin and anti-HA.

Fresh lysate containing fluorescently labeled receptors were then kept on ice for up to 2 hours until they were diluted using buffer containing 0.05% LMNG and 0.005% CHS in EX buffer and added to the flow chamber. When a desired single-molecule spot density (∼0.2 spots/μm^2^) was obtained, unbound receptors were washed with the dilution buffer. Single-molecule movies were recorded using a 100× oil immersion objective (NA 1.49) on an inverted microscope (Olympus IX83) with TIR mode at 20 Hz with 50-ms exposure time using two scientific sCMOS cameras (same as above, Hamamatsu ORCA-Flash4v3.0). Movies were recorded until >90% of molecules were bleached in the field of each movie. Data were analyzed using a custom-built LabVIEW (National Instruments) program^124^.

Briefly, movies from both channels were concatenated and loaded on the analysis program to visualize each channel for identification of colocalized molecules. Bleaching steps were assigned by inspecting the fluorescence traces manually for each molecule and plotting to show the bleaching step distribution. As previously established, the subpopulation of molecules showing 2 bleaching steps is a function of incomplete fluorophore labeling, incidental bleaching, and other detection issues with 45-55% 2-step bleaching typically seen for strict dimers^55, 81^. Colocalization and bleaching step analyses of GPCR-Barr complexes are interpreted as estimates on complex stability and are most likely lower than actual percentages in cells. This is because of expected complex dissociation during both cell lysis and upon immobilization of complexes on glass slides where dilution of the sample would decrease the total number and stoichiometries of complexes due to mass action. However, dimeric mGluRs are stable throughout the experiment due to their covalent attachment via disulfide bonds. Additionally, we observe little to no active dissociation of complexes when imaging complexes (which would be seen as increased background fluorescence in the arrestin channel). Under the assumption that all GPCR-arrestin complexes will be affected equally by these conditions, we expect similar differences in colocalization percentages will be retained in cells while the actual percentage may differ slightly. For photobleaching analyses, we also assume that values for 1 and 2 step photobleaching reported here are under-estimates of the true percentages of 2:1 and 2:2 mGluR: Barr complexes in cells.

### Single Molecule FRET

48 hours post-transfection with a modified SNAP-mGluR8 clone containing a C-terminal HA tag alone or in combination with GRK2 and β-arr1βCTD, HEK 293T cells were washed with EX buffer, and then labeled for 45 min at 37° C. with 1 μM of BG-LD555 (donor) and 3 μM of BG-LD655 (acceptor). Labeling solution was removed, cells were washed with EX buffer, and lysates were prepared as described above for SiMPull experiments. After immobilization of receptors, buffer was exchanged with imaging buffer and oxygen scavenging system with or without 1 mM glutamate as previously described^87^. FRET measurements were conducted using an Olympus IX83 microscope in TIR mode, using a 1.49 NA 100x oil immersion objective, and solid-state 561-nm laser for excitation. Fluorescence emission was filtered through a 635LP and detected using two synchronized sCMOS cameras (Hamamatsu ORCA-Flash 4v3.0). smFRET time lapses were captured at 30 ms exposure. FRET movies were analyzed using the program SPARTAN ^125^. Briefly, FRET efficiency was calculated using the following equation: 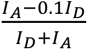 where 𝐼*_D_* is the donor intensity and 𝐼*_A_* is the acceptor intensity after subtracting background fluorescence. Molecules were selected for analysis if they exhibited canonical smFRET characteristics (e.g. inversely correlated donor and acceptor intensities; single bleaching steps in donor and acceptor channels) and were not selected based on specific fluorescence intensities corresponding to known LBD states (open vs. closed). Hidden Markov analysis was performed on the FRET traces using 3 states (0.15, 0.25, 0.35) to derive transition rates in SPARTAN. FRET histograms were plotted using GraphPad Prism and Gaussian fitting was performed using Origin.

### mGluR8 Purification and Complex Formation

Expi GNTi-cells (Thermo Fisher Scientific) were grown at 37 °C and 8% CO_2_ to a density of 3 × 10^6^ cells/mL in Expi media (Thermo Fisher Scientific) supplemented with 1:500 penicillin/streptomycin (Gibco) prior to transfection. Following manufacturer provided protocols, HA-FLAG-mGluR8 construct was transfected using the Expi293 transfection kit. The suspension was incubated at 37 °C for 20 h, then Expi enhancers were added to the cells as specified by the manufacturer (Thermo Fisher). Cell pellets were collected 72 hours after transfection by centrifugation at 4,000 rpm, flash-frozen in liquid nitrogen, and stored at −80 °C. For the mGluR8/β-arr1 complex, the steps above were followed except for co-transfecting HA-FLAG-mGluR8 with β-arr1-ΔCTD and GRK2-CaaX at a 2:1:1 ratio and adding 30 𝜇M L-AP-4 (Tocris)and 10 𝜇M VU6005649 (Tocris) 1 hour prior to harvesting cells. The same protocol was adopted for mGluR8/β-arr2 complexes using β-arr2-3A-ΔCTD instead of β-arr1-ΔCTD.

Cell pellets were Dounce homogenized in lysis buffer containing 20 mM HEPES (pH 7.5), 150 mM NaCl, 2 mM CaCl2, 30 𝜇M L-AP-4, and 10 𝜇M VU6005649. For mGluR8-β-arr1ΔCTD purifications, lysis buffer was supplemented with 3uM 8:0 PI(4,5)P2 (Avanti Polar Lipids). Protein was extracted via the addition of n-Dodecyl-β-D-maltopyranoside (DDM) with cholesteryl hemisuccinate (CHS) (Anatrace, D310-CH210) to a final concentration of 50 mM DDM/4 mM CHS and nutated for 1 hour at 4 °C. Following ultracentrifugation at 4° C at 96,000 × *g* for 30 minutes, the supernatant was filtered using a 0.22µm filter (VWR). A column containing 5 mL FLAG resin (Genscript) was then equilibrated with running buffer containing 100 mM HEPES (pH 7.5), 150 mM NaCl, 2 mM CaCl_2_, 0.1% glyco-diosgenin (GDN), and 30 𝜇M L-AP-4. Supernatant was then added to FLAG column resin and incubated at room temperature for 1 hour while being stirred. Resin was then reapplied to the column and flow through was collected. The column was washed first with 10 mL running buffer supplemented with 10 mM MgCl_2_ and 5 mM ATP to remove contaminating heat-shock proteins, followed 10 mL of running buffer. This series of two washes was then repeated to ensure removal of any contaminants and to facilitate detergent exchange. FLAG-mGluR8 was subsequently eluted with running buffer supplemented with 0.2 mg/mL FLAG (DYKDDDDK) peptide (GenScript). Elution fractions were collected, analyzed by SDS-PAGE, and concentrated using a 50 kDa molecular weight cut-off spin concentrators (Amicon). The sample was then loaded onto a Superose 6 Increase 10/300 GL (Cytiva) column equilibrated in SEC buffer containing 100 mM HEPES (pH 7.5), 150 mM NaCl, 2 mM CaCl_2_, 0.01% GDN, and 30 𝜇M L-AP-4. Elution fractions were collected and analyzed by SDS-PAGE for purity. Key fractions were then concentrated using a 50 kDa molecular weight cut off spin concentrator.

For purifications containing mGluR8/β-arr1-ΔCTD complexes, protein was incubated with 2.5 µM Fab7 for 1 hour at room temperature, followed by addition of Sulfo-LC-SDA (Thermo Scientific) crosslinker to a final concentration of 250 µM. Samples were then incubated for 45 minutes at room temperature in the dark. To quench NHS-crosslinking Tris was added to a final concentration of 3 mM and samples were then irradiated with 365 nm light using an ENF-240C UV lamp (Spectro-UV, 4 watt tube) for 45 minutes at 4 °C. To increase arrestin association, 1uM 8:0 PIP2 was added and allowed to incubate overnight at 4 °C. Crosslinked samples were then purified using a second anti-FLAG column following the same protocol and using the same buffers as described above. Final samples for both mGluR8 and mGluR8/β-arr1ΔCTD complexes were concentrated to 3-5 mg/mL using a 50 kDa molecular weight cut off spin concentrator (as above) and used for negative stain and cryo-EM studies. Protein was either used immediately or flash frozen in liquid nitrogen and stored at -80 °C. For purifications containing mGluR8/β-arr2-3A-ΔCTD complexes, protein was subjected to a second anti-FLAG column similar to mGluR8/β-arr1-ΔCTD complexes, but did not receive additions of crosslinker, Fab7, or PIP2.

For mGluR8/G protein complexes, purified FLAG-mGluR8 (as described above) was added to purified G protein heterotrimers containing G_⍺i1_, G_β1_, and G_𝛄2_ (provided by Kaavya Krishna Kumar) at a final molar ratio of 1:3 in a total reaction volume of ∼100 µL. The mixture was incubated for 3 hours at room temperature in an Eppendorf tube, followed by the addition of 2 µL of apyrase (100 units/mL from New England BioLabs) and a further incubation for 1.5 hours on ice. The single-chain antibody fragment scFv16 was added to stabilize the mGluR8/G protein complex at a 1.5:1 molar ratio relative to receptor and incubated overnight at 4 °C. The complex was then purified using an anti-FLAG resin column (GenScript) as described above, in the presence of both 30 µM L-AP4 and 10 µM VU600. Purified mGluR8-G_i1_ complexes were collected and concentrated as described above in the mGluR8-arrestin purification procedure.

### Mass Spectrometry

Purified mGluR8/β-arr1 preparations were dissolved in SDS lysis buffer (10 mM (tris(2-carboxyethyl)phosphine) (TCEP), 2% SDS, 150 mM NaCl, 50 mM Tris, pH 8.5) and reduced cysteine residues were alkylated with iodoacetamide at a final concentration of 20 mM and incubation at room temperature in the dark for 25 minutes. Alkylation was quenched with 50 mM freshly prepared DTT. SDS was removed by SP3 protein cleanup^126^ and proteins were digested with Lys-C protease (Wako, Catalog Number 129–02541) and trypsin (Promega Catalog Number V5113). Peptides were subjected to micro-phospho enrichment as described recently in Susa et al., 2024^127^ and analyzed on an Orbitrap Ascend Mass spectrometer (Thermo Fisher Scientific) using 180 or 300 min FAIMS runs (CVs -40V, -55V, -70V, -85V, cycle time 6s) at an MS1 resolution of 120,000 in the orbitrap. After HCD fragmentation (25% collision energy), MS2 scans were collected at 45,000 resolution in the orbitrap with a 1.6 m/z quadrupole isolation window. Data were searched with COMET^128^ using additional masses of +15.994914 Da for oxidized methionine modification and + 79.966330 Da for phospho-modification on S, T and Y residues, dynamically. Peptides were identified using MS2 spectra and a false discovery rate (FDR) < 1% and was achieved by applying a target-decoy database search strategy. Details of phospho-proteomic analysis are provided in Susa et al., 2024^127^.

### Negative Stain EM

An aliquot of 3.5 µL of mGluR8 or mGluR8/β-arr1-ΔCTD or mGluR8/β-arr2-3A-ΔCTD samples was applied onto glow-discharged carbon-coated grids (Electron Microscopy Services, Cu 400 mesh) and allowed to adsorb for 30 s. Excess sample was blotted off using filter paper. Then 5 µL of freshly prepared 0.75% (w/v) uranyl formate stain was added for 30 s and immediately blotted. Another 5 µL of stain was then added to the grid for 5 minutes, followed by blotting and benchtop air-drying for 5 minutes.

Grids were imaged on a Talos L120C electron microscope (FEI) equipped with a 4K x 4K OneView CMOS camera (Gatan, Inc.) at a nominal magnification of 92,000x, corresponding to a pixel size of 1.6 Å. Manual data collection was performed with a defocus ranging from -0.4 µm to -0.8 µm and a dose of approximately 60 electrons per Å^2^ with 1 s of exposure time. CryoSPARC was used to collect and process micrographs, manually pick particles, and perform reference-free 2D classification.

### Cryo-EM

mGluR8 or mGluR8/β-arr1-ΔCTD samples ranging from 3-5 mg/mL were applied at a volume of 3 µl to plasma treated UltrAuFoil 1.2/1.3 300 mesh grids (Quantifoil). Vitrified samples were prepared using a Vitrobot Mk IV (Thermo Fisher), blot time of 2 seconds, blot force of -2, at 100% humidity and 22^°^C. Grids were imaged with a Titan Krios electron microscope (Thermo Fisher) operated at 300 kV and a nominal magnification of 105,000× and equipped with a K3 camera (Gatan) set in super-resolution mode (0.426 Å pixel size). Images were collected using Leginon.

For mGluR8 alone datasets, beam induced motion was corrected using MotionCor2 in Relion 3.1 using two-fold binning and dose-weighting. These images were then used for contrast transfer function estimation via CTFFIND4.1^129^. In Relion, particles were picked using a 3D initial model template of the receptor and extracted using a box size of 416. Further processing was then performed in CryoSPARC version 3.3.2^130^. Ab initio followed by multiple rounds of heterogeneous refinement resulted in two good classes that were further refined via subsequent ab initio and heterogeneous refinement. The best heterogeneous refinement class/classes were then used for homogeneous refinement followed by non-uniform refinement with C1 symmetry. These particles were then imported back into Relion for Bayesian polishing^131^ followed by non-uniform refinement to obtain the best full-length map. Initial mGluR8 models were built using AlphaFold2^132^. Each model was then refined via iterative rounds of real-space refinement using the full-length map in Phenix^133^ and manual refinement in Coot^134^. Regions with poor resolution were either deleted, as in the case of some loops, or stubbed, as is the case of the TMDs. Visual representations were prepared using ChimeraX^135^ and Pymol (Schrödinger, LLC 2015).

For datasets containing the mGluR8/β-arr1-ΔCTD complex, micrographs were motion corrected using Warp^136^ and imported to CryoSPARC v4.5.3 for processing. Contrast transfer function estimation was performed using the PatchCTF job and particles were picked using an initial 3D model template of the complex and extracted with a box size of 588. Four iterative rounds of 2D classification were followed by ab initio and heterogeneous refinement, further 2D classification, and a second round of ab initio and heterogeneous refinement revealed one promising class of particles. This class was then split into two further classes following ab initio, heterogeneous, and non-uniform refinement with C1 symmetry to obtain best full-length maps. These maps contained higher resolution density for mGluR8 and lower resolution density that was assigned to β-arr1ΔCTD. An initial model of the mGluR8/β-arr1 complex was also created from this density using our mGluR8 preactive class 1 structure as a starting point. The model of mGluR8 was refined in Phenix and Coot as described above. To place the β-arr1ΔCTD protomer, AlphaFold2 (AF2) multimer^137^ was used to generate complexes of mGluR8 TMD dimers with a single copy of β-arr1ΔCTD. The CTDs of mGluR8 were removed for AF2 modeling due to their predicted disorder, the potential presence of “conditionally folded” regions of the CTD^138^, and the inability of AF2 to phosphorylate residues. We found that AF2 produced a model that fit our density well, and this model was used to initially place the β-arr1 near our full length mGluR8 model. Further rigid-body refinement of the β-arr1 into our density was performed using Coot.

The model containing the most β-arr1 density was then used as a template to pick particles using Cryosparc template picker. Subsequent templated heterogeneous refinement using 3 starting models: preactive mGluR8 class 1, our initial mGluR8/β-arr1 model, and a junk class. This resulted in a class with a much more defined β-arr1 density with clear domain orientation. Further local refinement was performed on the mGluR8 density to better resolve receptor features in the presence of β-arr1 (Fig. S11). A final mGluR8/β-arr1 model was then constructed using our initial model as a template. Regions with poor resolution were either deleted, as in the case of some loops, or stubbed, as is the case of the TMDs. Visual representations were prepared using ChimeraX^135^ and Pymol (Schrödinger, LLC 2015). Structural comparison and alignment with existing GPCR-β-arr1 structures were performed in Chimera X by aligning the receptor using the ‘matchmaker’ function. Angles were measured as the difference in the middle reference plane change between arrestins (Fig. 5A) or from receptor to arrestin (Fig. 5B).

For mGluR8/G protein complexes, samples were frozen on gold grids using the same procedures as described above for other mGluR8 cryo-EM samples. Cryo-EM data were auto-collected at the Weill Cornell Medicine Cryo-EM facility in two batches using EPU on a 200 kV Glacios transmission electron microscope (Thermo Fisher Scientific) equipped with a Falcon 4i direct electron detector. The microscope was operated with a 2.7 mm spherical aberration objective lens and a 100 µm objective aperture. Movies were recorded at a nominal magnification of 100,000×, corresponding to a calibrated pixel size of 1.16 Å, with a defocus range of –1.8 to –0.8 µm. Data were acquired in electron event representation (EER) mode with a total exposure time of 4.69 s, corresponding to an accumulated dose of 30 e⁻/Å². An energy filter with a 10 eV slit width was applied during data collection.

All image processing was performed in cryoSPARC (v.3.2.2). After patch-based motion correction and CTF estimation, micrographs were curated based on CTF fits prior to further processing. An initial particle set was selected from the first batch of micrographs using template-based picking with projections of an AlphaFold3-generated^139^ mGluR8–G_i1_ complex model. Template-picked particles were subjected to three rounds of two-dimensional classification and heterogeneous refinement using templates to sort particles into three classes: mGluR8–G-protein (described above), mGluR8 alone (templated from mGluR8 class 1 structure), and a junk class. Heterogeneous refinement resulted in a class of approximately 250,000 particles corresponding to the mGluR8-Gprotein complex, which was refined by non-uniform refinement.

A 50,000-particle subset of the final heterogenous class described above was used to train the Topaz neural network-based particle picker^140^. The trained model was then applied to the full curated dataset, and extracted Topaz-picked particles were further assessed by two-dimensional classification (2DC). Picked particles from the 2DC were subjected to heterogeneous refinement and 3D classification, focusing on the mGluR8–G-protein class. Particles from the best-resolved classes underwent *ab initio* reconstruction followed by heterogeneous refinement without templates. The final subset of particles was refined using non-uniform refinement and subjected to local refinement. Masks were generated using ChimeraX and the Volume Tools module in cryoSPARC with soft masks applied to individual regions, including the LBDs, CRD + TMDs, TMDs + G proteins, and the G proteins alone. To generate a composite reconstruction, density maps from individual refinements were aligned to the global map in UCSF ChimeraX^135^ and combined using the *vop maximum* command.

### Molecular Dynamics Simulations

Atomistic molecular dynamics (MD) simulations were performed on the full-length mGluR8/β-arr1 complex embedded in a lipid bilayer with composition mimicking that of the plasma membrane. To build this full-length, lipid embedded mGluR8/βarr1 complex, we combined the mGluR8/β-arr1-ΔCTD model derived from the combination of cryo-EM experiments with AlphaFold2 predictions and rigid body docking as described above. To mimic the biologically relevant system, we docked a peptide containing the fully phosphorylated mGluR8 CTD ST-rich region (residues 888 to 908) to the β-arr1 in the same orientation as a V2R-CTD/β-arr1 cryo EM structure (PDB ID: 4JQI). Additionally, we replaced the orthosteric agonist L-AP-4 with glutamate and removed the PAM (VU600) in our model to mimic the endogenous form of the mGluR8/β-arr1 complex. Using CHARMM-GUI membrane builder^141^, the protein complex was embedded in a lipid bilayer containing POPC, POPE, POPS, cholesterol, sphingomyelin, and PIP_2_ at approximately biological amounts^142^. There were 20 cholesterol, 196 POPC, 20 POPE, 16 POPS, and 200 sphingomyelin molecules in the upper leaflet and 16 cholesterol, 104 POPC, 172 POPE, 116 POPS, 32 PIP_2_ molecules in the lower leaflet. The protein-membrane system was then solvated and ionized in CHARMM-GUI using 0.2 M NaCl concentration. Overall, the system contained 792,623 total atoms including water, sodium, and chloride ions. Residues were protonated as recommended by CHARMM-GUI to reflect pH 7.0 (all histidine residues were deprotonated except residue 55).

Using NAMD version 2.14, the initial system was separately equilibrated three times (i.e. three independent replicates) using the standard CHARMM-GUI equilibration protocol. Each of the resulting three equilibrated systems were then used to seed a new set of 24 ∼267 ns-long independent simulations (with randomly resetting velocities of all the atoms in the system). Thus, 72 independent replicates were collected with the cumulative MD sampling time of ∼19 μs. These production runs were carried out with OpenMM version 7.5.1 using the following run parameters: 4 fs integration timestep enabled with hydrogen mass repartitioning, Particle Mesh Ewald (PME) for electrostatic interactions, under isothermal-isobaric (NPT) ensemble conditions using semi-isotropic pressure coupling, with MonteCarloMembraneBarostat and Langevin thermostats used to maintain constant pressure and temperature, respectively. Additional simulations parameters included: “friction” set to 1.0/picosecond, “EwaldErrorTolerance” 0.0005, “rigidwater” True, and “ConstraintTolerance” 0.000001. The van der Waals interactions were calculated applying a cut-off distance of 12 Å and switching the potential from 10 Å.

Simulations were analyzed using VMD plugins and home-made TCL and python scripts. Statistical tests were performed using GraphPad Prism10. All analyses were performed on the trajectories strided with a timestep of 0.4 ns, except the buried SASA calculation which was calculated at a timestep of 0.8 ns (sampled half as often as the other analyses for computational efficiency). To determine the x-y angles of β-arr1 relative to mGluR8, the principal axes of mGluR8 were determined using the Orient plugin in VMD and compared to a vector connecting the center-of-mass of the β-arr1 N and C lobes. To determine the β-arr1 z-angle, we compared the same β-arr1 vector to a vector along the membrane normal. The x-y/z angle heatmap was created by binning corresponding x-y and z angles for β-arr1 in each frame of each trajectory in the 72 replicas in 5 degree increments. RMSD values were calculated by aligning individual protomer chains and using the VMD RMSD command. Residue-to-residue and domain center-of-mass distances were calculated by measuring the length of the vector connecting the two points. Lipid hydrogen bond analyses were carried out using the default settings in the VMD hydrogen bond calculator using default settings for bond lengths and angles and buried SASA analyses were calculated using the VMD “sasa” function using a probe radius of 1.4 Å as previously^70^. To create a contact heatmap for *cis* and *trans* contacts for β-arr1 with mGluR8, contacts were calculated by adapting code from the Klauda lab (https://user.eng.umd.edu/∼jbklauda/wiki/doku.php?id=protein_contact_map) to calculate contacts with a cutoff distance of 5Å for each residue in the mGluR8 TMD with each residue of β-arr1. These resulting contacts for each residue were then scaled relative to the total frames in the simulation to get a value for contact frequency.

### Single cell RNA sequencing analysis

We analyzed previously obtained single cell RNA sequencing data from the Anterior Cingulate Cortex (ACA SMART-seq, Allen Institute, 2018)^103^. All cells passed the quality control criteria and underwent subsequent hierarchical clustering according to the similarity of their individual transcriptomes^143^. A total of 5028 cells included from the mouse ACC cortex were grouped by cell type according to the different brain layers and populations of interest (L2/3, L2/5, L5, L5/6, L6, L6b glutamatergic and NGF, Sst+, PV+ and VIP GABAergic neurons). For co-expression analysis of *Arrb1* and *Arrb2* (Fig. 7 and S9), we took into consideration the low rate of false positives and under-sampling of a cell’s total mRNA with this technique, thus imposing a threshold of five copies per million (CPM). Data were analyzed using Microsoft Excel and R Studio for plotting (*ggplot* package).

## Acknowledgments

We thank the entire Levitz lab (Weill Cornell Medicine), Jonathan Javitch (Columbia University), and Kaavya Krishna Kumar (Stanford University) for helpful discussions. We also thank the Weill Cornell Medicine Cryo-EM Core facility, the Weill Cornell Scientific Computing Unit, the NYU Langone Health Microscopy Laboratory, and the NYU Langone Health Cryo–EM Laboratory. This work is supported by NIH grants F31NS129320 (A.S.), F32GM148001 (D.M.), R01NS129904 (J. Levitz), the Margarita Salas Fellowship from the Ministry of Universities of Spain (A.G.-H.), the Charles Revson Fellowship (A.G.- H.), the Rohr Family Research Scholar Award (J. Levitz), and the Monique Weill-Caulier Award (J. Levitz).

