## SupplementalFigures for "Structural Diversity and Dynamics of Metabotropic Glutamate Receptor/Beta-Arrestin Coupling"

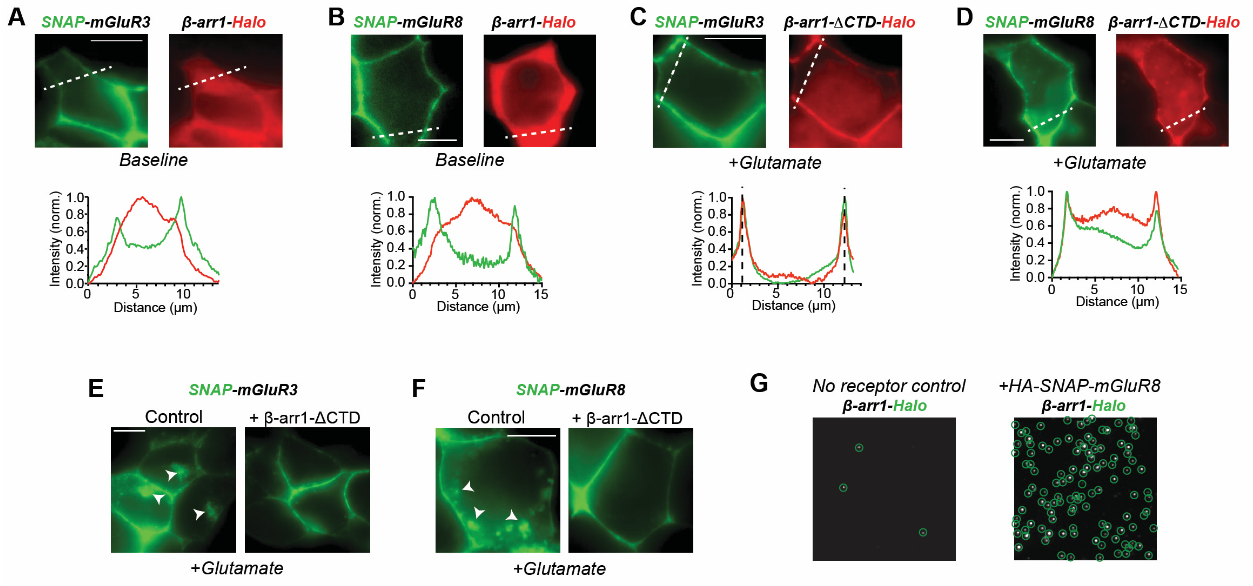


**Figure S1. Further analysis of β-arr coupling of mGluR3 and mGluR8. (A-B)** Representative images showing lack of surface localization of β-arr1 with mGluR3 (A) or mGluR8 (B) in the absence of glutamate treatment. **(C-D)** Representative images showing surface localization, but limited co-internalization, of β-arr1ΔCTD with mGluR3 (C) or mGluR8 (D) in the presence of glutamate treatment. **(E-F)** Representative images showing decreased internalization of mGluR3 (E) and mGluR8 (F) upon co-expression of β-arr1ΔCTD (right). **(G)** Representative images showing mGluR-dependence of β-arr1-Halo immobilization in SiMPull. Green circles indicate immobilized spots. Scale bar: 5 µm.

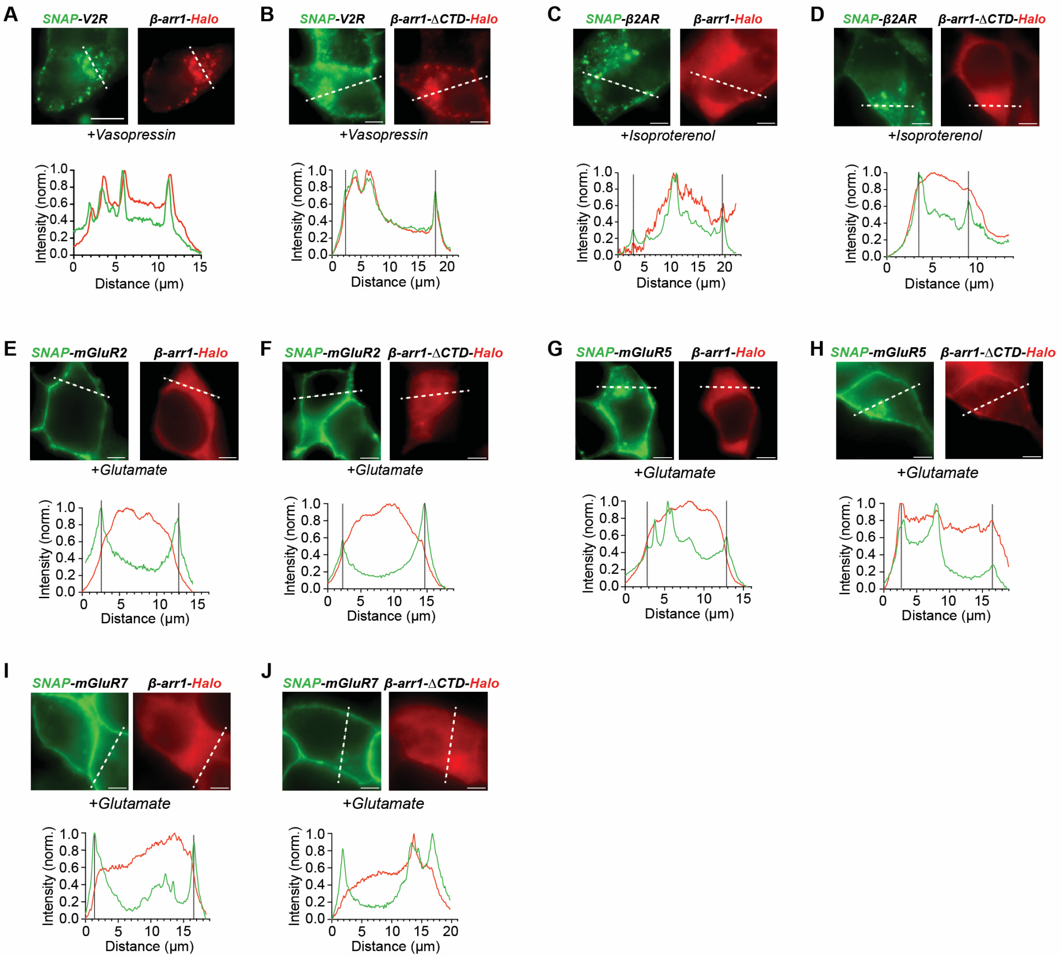


**Fig. S2. Further cell imaging of β-arr1 coupling to a panel of GPCRs. (A-J)** Representative images and line scans showing surface localization and/or co-internalization of β-arr1 or β-arr1ΔCTD with a range of GPCRs following agonist treatment for 30 min. Scale bar: 5 µm.

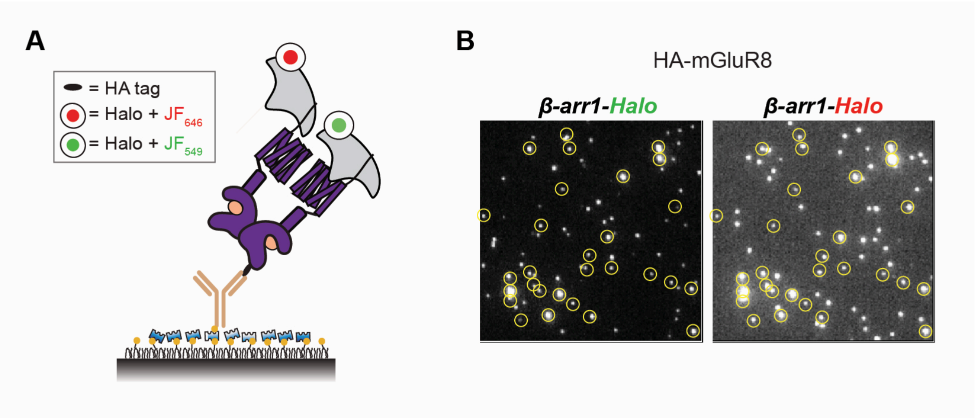


**Fig. S3. Further SiMPull analysis of GPCR/β-arr1 coupling stoichiometry. (A-B)** Schematic (A) and representative images (B) showing co-localization of β-arr1-Halo labeled in two different colors upon co-pulldown with HA-mGluR8. Yellow circles (B) indicate colocalized spots indicative of mGluR/ β-arr1 complexes containing more than one β-arr1.

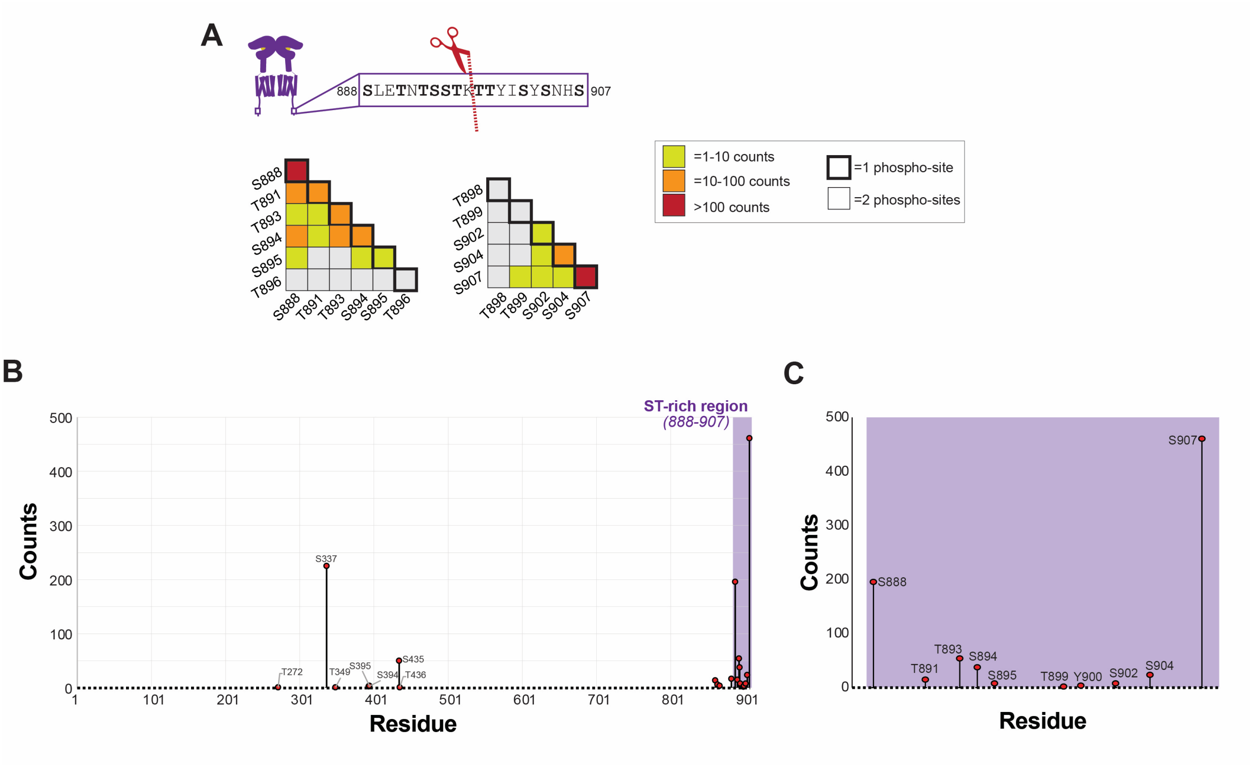


**Fig. S4. Mass spectrometry analysis of mGluR8 phosphorylation. (A)** Top, sequence of distal region of mGluR8 CTD showing protease cut site (red). Bottom, summary of mass spectrometry analysis of phosphorylation within each resulting peptide. **(B-C)** Mass spectrometry analysis of mGluR8 showing abundance of detected phosphosites with strong enrichment in the distal C-terminus.

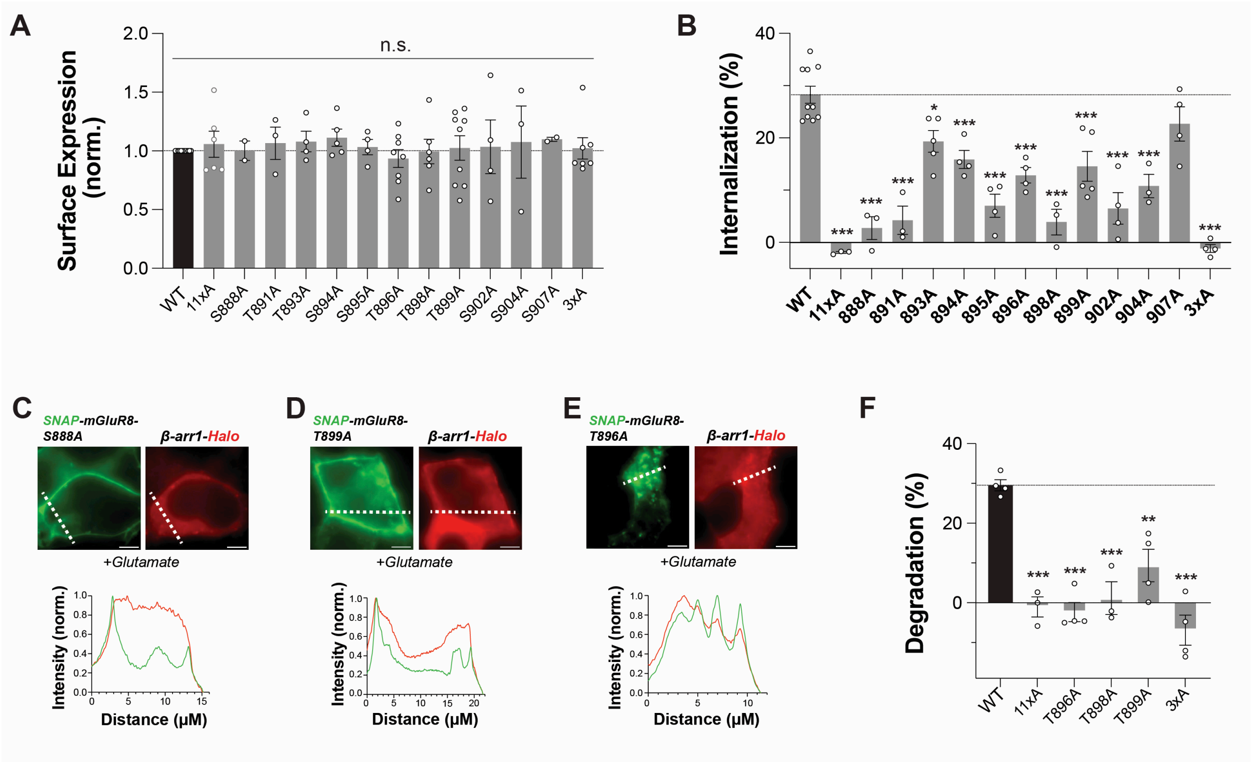


**Fig. S5. Further analysis of mGluR8/β-arr tail coupling. (A)** Quantification of surface expression levels for all mGluR8 CTD mutations. **(B)** Quantification of degree of mGluR8 internalization for all CTD mutants showing a significant decrease relative to WT for most mutations. **(C-E)** Representative images and line scans showing effects of CTD mutants on mGluR8 internalization and β-arr1 localization. **(F)** Quantification of degree of mGluR8 degradation for a subset of CTD mutants. Points represent individual experiments (A, B). 1-way ANOVA with Dunnet's multiple comparisons test (to WT) is used in (A, B, F). *, p<0.5; **, p<0.01; ***, p<0.001. All data shown as mean ± SEM. Scale bar: 5 μm.

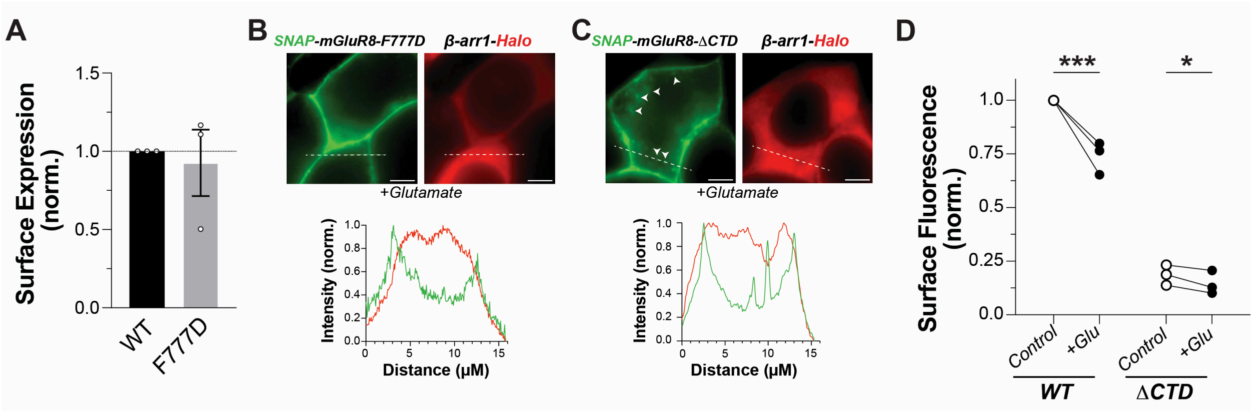


**Fig. S6. Further analysis of mGluR8/β-arr core coupling. (A)** Surface expression analysis showing a lack of effect of introduction the F777D mutation into mGluR8. **(B)** Representative image and line scan showing lack of internalization and β-arr recruitment by mGluR8-F777D. **(C)** Representative image and line scan showing modest internalization of mGluR8-ΔCTD. Arrows highlight intracellular receptor puncta. **(D)** Summary graph showing that mGluR8-ΔCTD shows substantially less surface expression compared to WT, but still undergoes a small glutamate-induced reduction in surface levels. Points represent individual experiments (D). Paired t-test is used in (D). *, p<0.5; ***, p<0.001. All data shown as mean ± SEM. Scale bar: 5 µm.

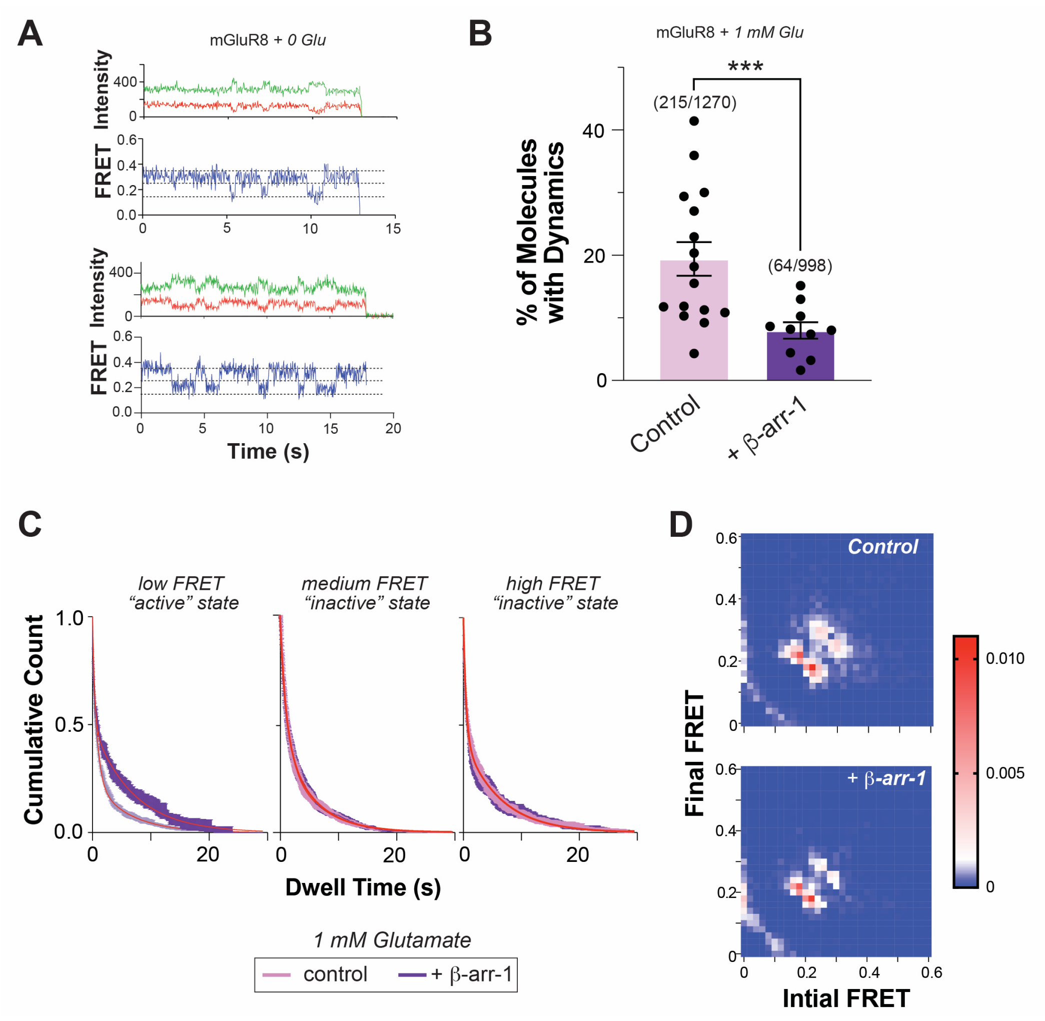


**Fig. S7. Further smFRET analysis of conformational effects of β-arr1 on mGluR8. (A)** Representative smFRET traces showing basal dynamics of mGluR8 in the absence of ligand. **(B)** Summary graph showing a reduced proportion of mGluR8 molecules showing dynamic inter-state transitions in the presence of β-arr1. **(C)** Survival plots showing a lack of an effect of β-arr1 on time spent in the medium (0.25) or high (0.35) FRET states, but a clear effect on the time spent in the low (0.15) FRET state, in the presence of 1 mM glutamate. **(D)** Transition density plots showing frequency of transitions between FRET states in the absence (left) or presence of β-arr1 (right) in the presence of 1 mM glutamate. Points represent individual movies (B). Unpaired t-test was used in (B). ***, $p<0.001$. All data shown as mean ± SEM.

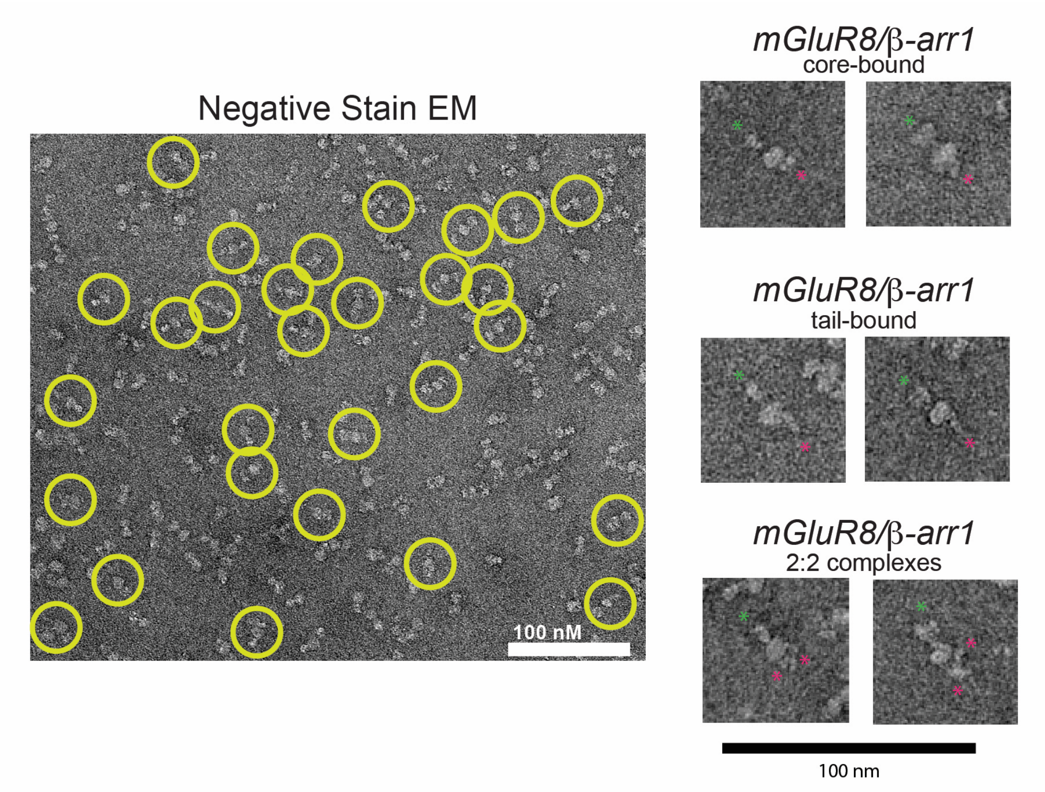


**Fig. S8. Representative mGluR8/β-arr1 negative stain particles.** EM micrograph showing identified mGluR8/β-arr1 complexes (yellow circles), left, and representative single particles for each major class of interactions, right. Red stars highlight β-arr1 molecules and green stars highlight mGluR8 LBDs.

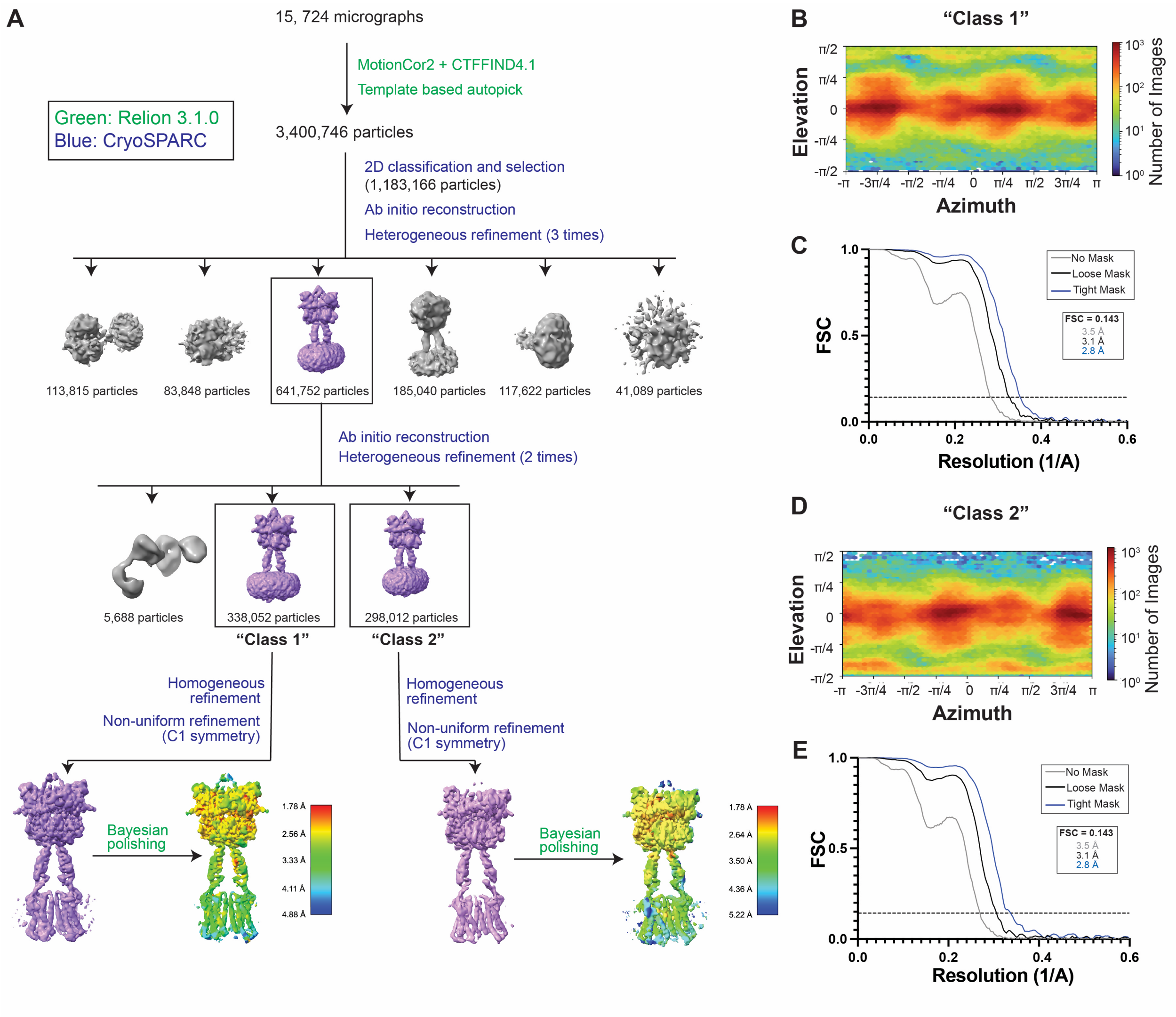


**Fig. S9. mGluR8 cryo-EM processing pipeline. (A)** Summary of processing pipeline for mGluR8 structures. **(B-E)** Angular distribution heatmaps and Fourier shell correlation (FSC) curves of particles in reconstructed mGluR8 maps.

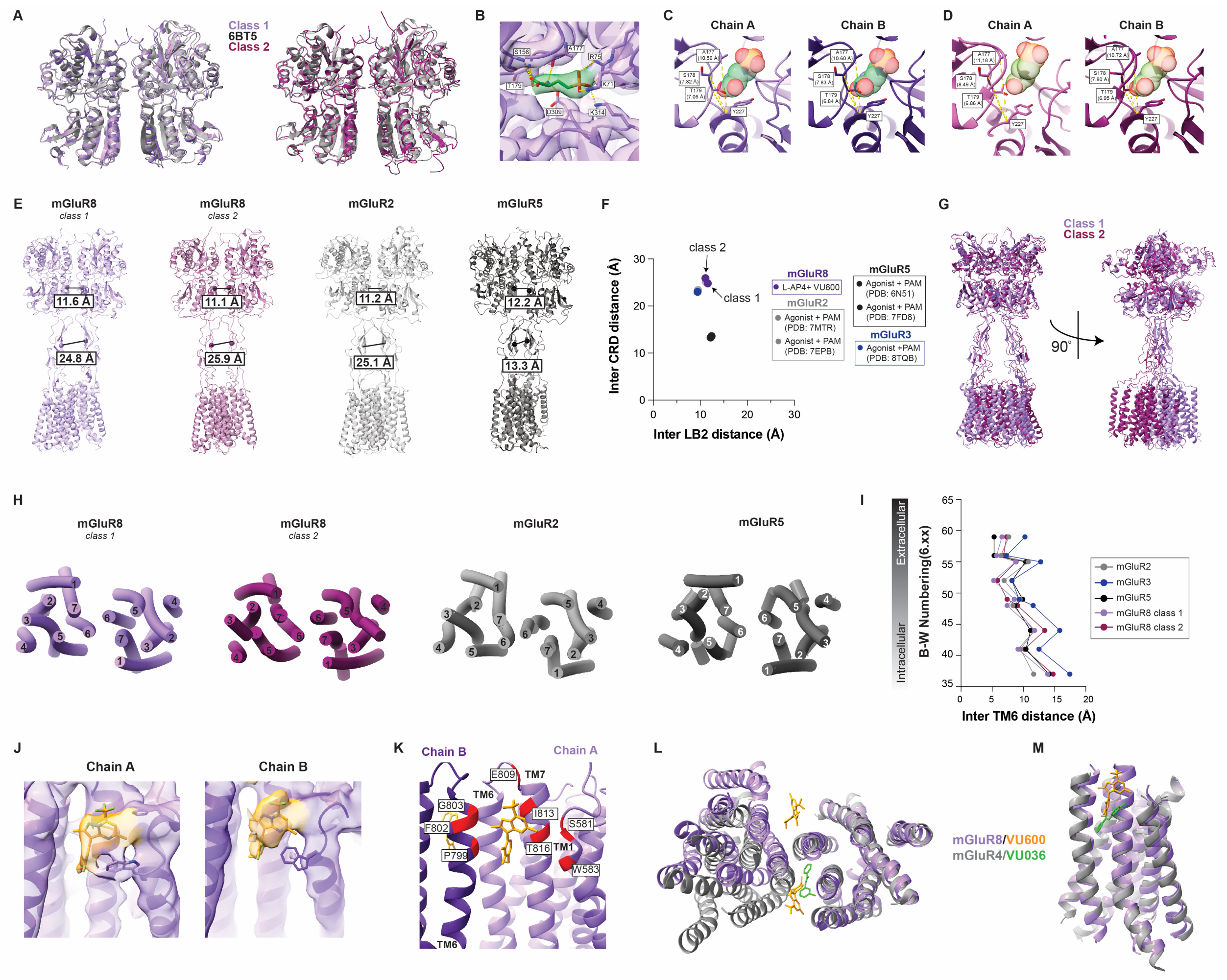


**Fig. S10. Structural analysis of mGluR8 cryo-EM structures. (A)** Structural alignment of class 1 and class 2 LBD dimers with a prior X-ray crystal structure (PDB:6BSZ), showing comparable degrees of LBD closure and an "active" lower lobe interface. **(B)** Zoom-in image showing density of L-AP4 in green with interactions with the ligand of distance less than 3.5 Å indicated. (S156-3.46 Å, T179 – 3.37 Å and 3.18 Å, D309-2.74 Å, K314-2.57 Å, K71-3.29 Å, R75- 2.30 Å and 2.51 Å, A177-2.34 Å. **(C, D)** Zoom-in showing modeled L-AP-4 binding position in chain A and chain B of class 1 (C) and class 2 (D) mGluR8 cryo-EM structures with measurements between top lobe residues and the bottom lobe residue indicated. **(E-F)** Comparison of mGluR8 cryo-EM structures with other agonist and PAM-bound mGluR structures showing a common tight LB2 (lower lobe) and inter-CRD interface. **(G)** Alignment of class 1 and class 2 mGluR8 structures at the LBD dimer reveals a large CRD/TMD offset as the major difference between classes. **(H-I)** Comparative analysis showing a common TM6-TM6 interface across agonist and PAM-bound mGluR structures. **(J)** Zoom-in showing density used to guide modeling of mGluR8 PAM VU600 in both chains. **(K)** Putative binding site of VU600 incorporating TM1, TM6 and TM7 residues from both chains. **(L-M)** Comparison of proposed PAM binding sites in mGluR and a prior G protein-bound mGluR4 structure (PDB: 8JD6) aligned to one protomer.

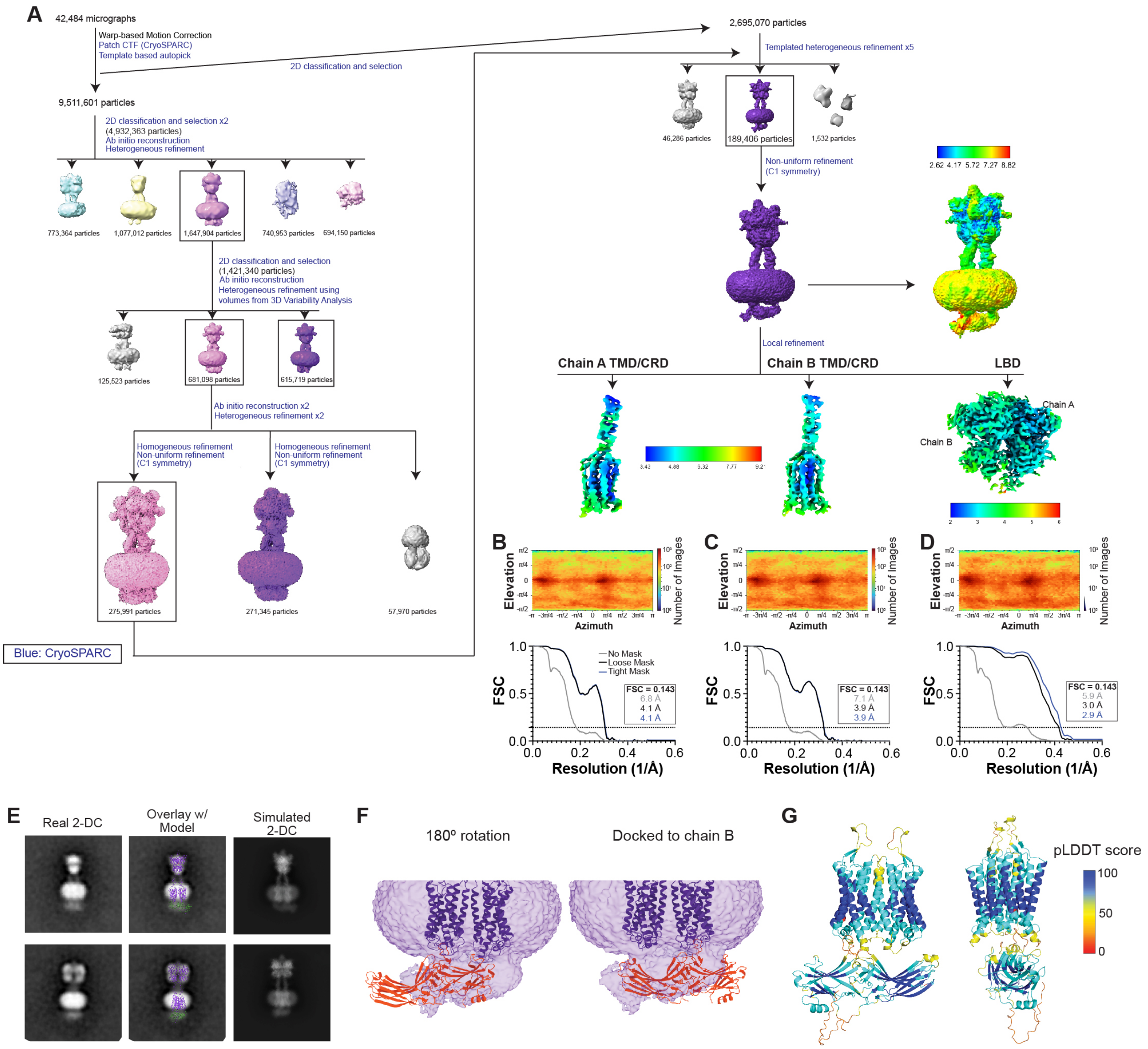


**Fig. S11. mGluR8/β-arr1 cryo-EM processing pipeline. (A)** Summary of processing pipeline for mGluR8/β-arr1 structures. **(B-D)** Angular distribution heatmaps and Fourier shell correlation (FSC) curves for locally refined domains in reconstructed mGluR8/β-arr1 maps. **(E)** Comparison of negative stain and cryo-EM data. "Real 2-DC" shows negative stain 2-D classes, "Overlay w/ model" shows the cryo-EM based model overlaid on the same 2-D classes, "Simualted 2-DC" shows the simulated 2-D classes calculated from the cryo-EM based model. **(F)** Image showing poor fit of β-arr1 in cryo-EM density if rotated 180 degrees or docked to the other TMD relative. **(G)** AlphaFold2 model of mGluR8 TMDs and β-arr1-ΔCTD which best matches cryo-EM density, colored by pLDDT scores.

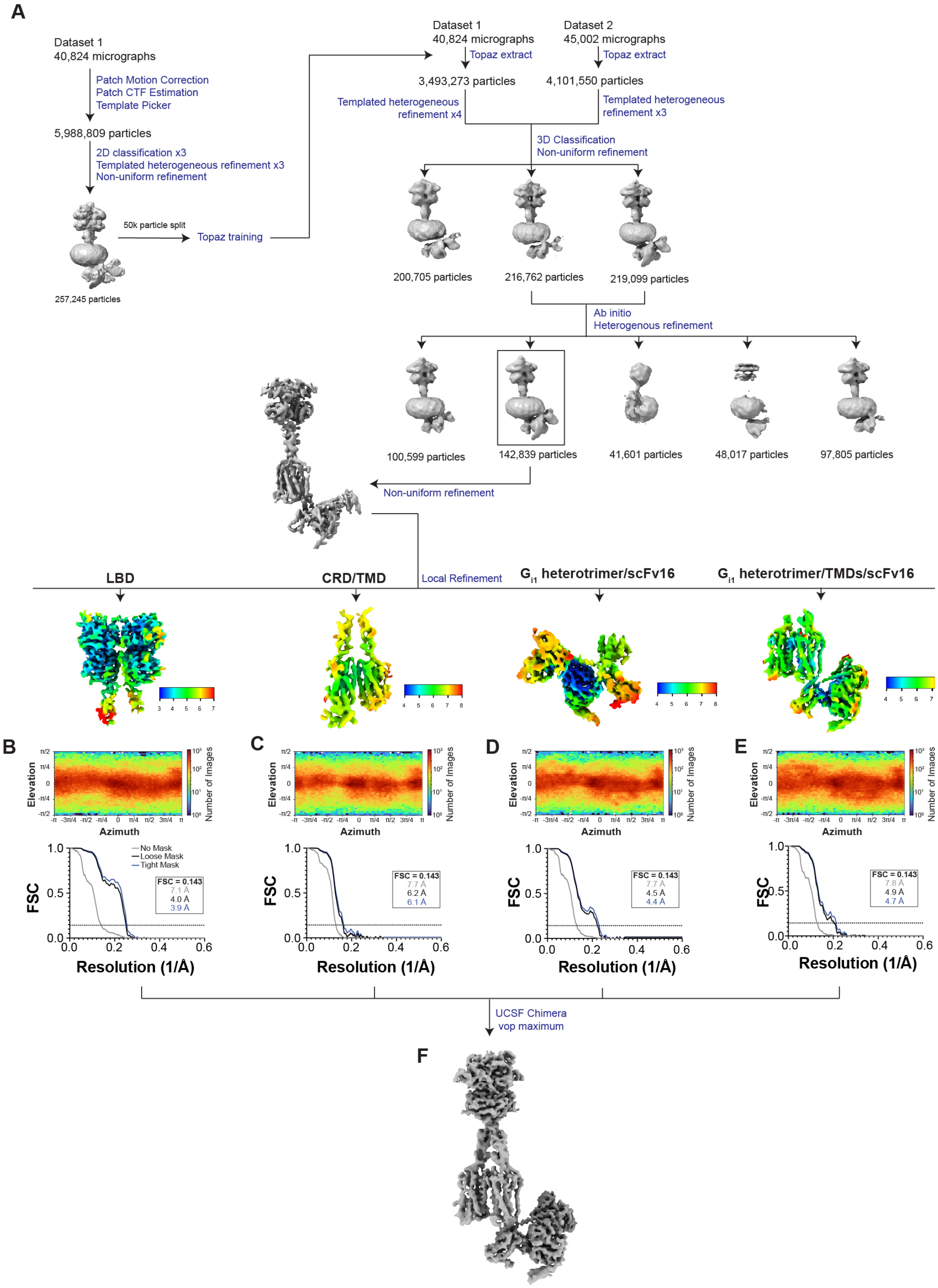


**Fig. S12. mGluR8/G protein cryo-EM processing pipeline. (A)** Summary of processing pipeline for mGluR8/G protein structure with **(B-E)** Angular distribution heatmaps and Fourier shell correlation (FSC) curves for locally refined domains in reconstructed maps. **(F)** Composite map used for model building.

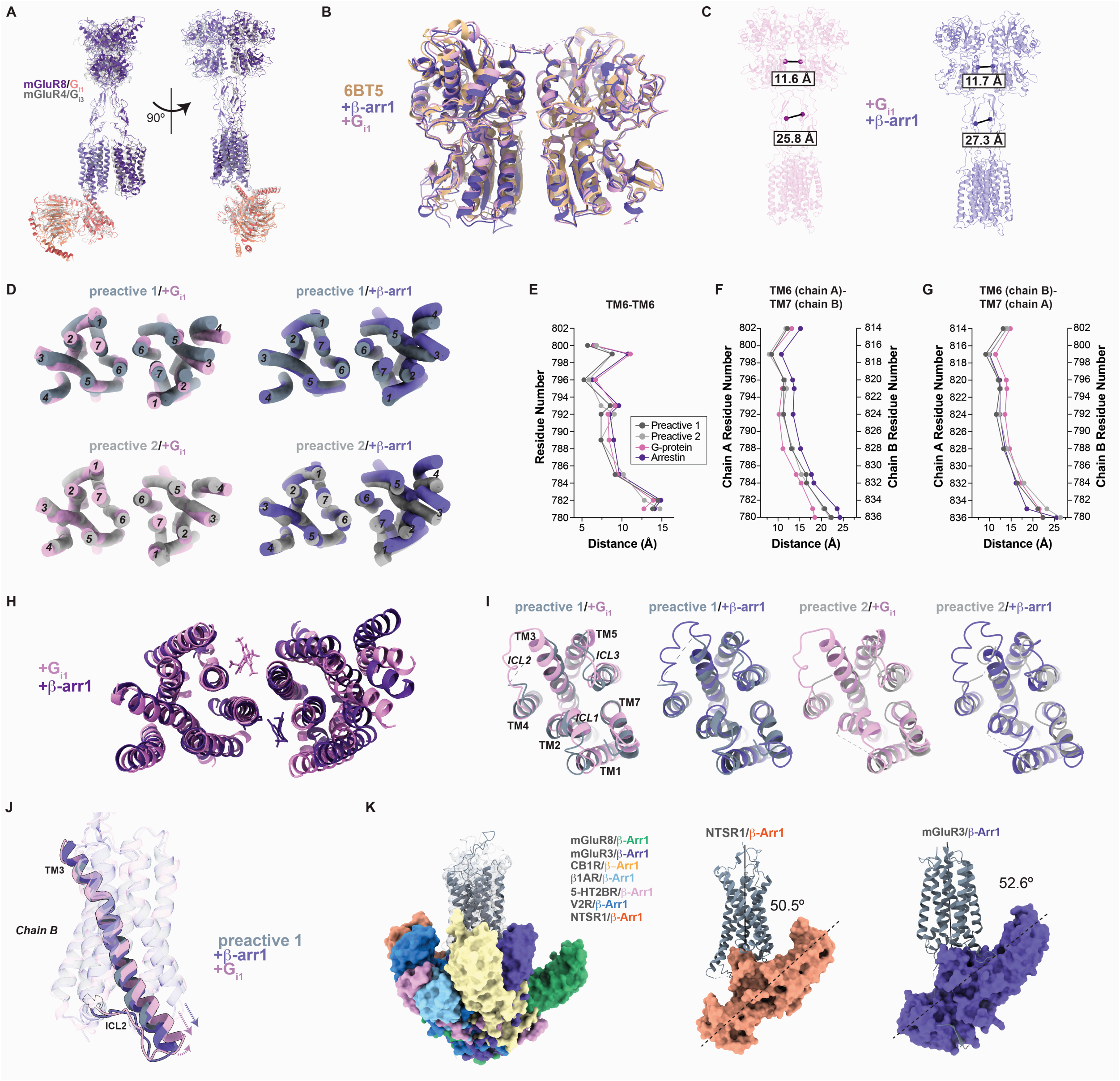

**Fig. S13. Further analysis of mGluR8/β-arr1 and mGluR8/G protein cryo-EM structures. (A)** Alignment of G protein-bound mGluR8 and mGluR4 structures showing a very similar overall arrangement. **(B)** Structural alignment of β-arr1 and G protein-bound mGluR8 LBD dimers with a prior X-ray crystal structure (PDB:6BSZ), showing comparable degrees of LBD closure and an "active" lower lobe interface. **(C)** β-arr1- and G protein-bound mGluR8 structures show a tight LB2 (lower lobe, E229) and inter-CRD interface (C553). **(D)** Top (extracellular) view showing slightly different TM6-TM6 interface between mGluR8 alone and β-arr1 and G protein-bound **(E-G)** Measurements of inter-TMD distances along TM6-TM6 (E), TM6 (chain A)-TM7 (chain B) (F), and TM6 (chain B)- TM7 (chain A) (G) interfaces, showing subtle differences across structures. **(H)** Top view showing distinct PAM locations for G protein and β-arr1-bound mGluR8 structures. **(I)** Bottom (intracellular) view showing subtle conformational rearrangements of chain A TMD across mGluR8 structures. **(J)** Alignment of chain B TMD showing TM3 elongation and lack of ICL2 repositioning in the presence of transducers. **(K)** Comparison of the degree of β-arr1 tilt (z-angle) in the mGluR8/β-arr1 model compared to a range of GPCR/β-arr1 structures.

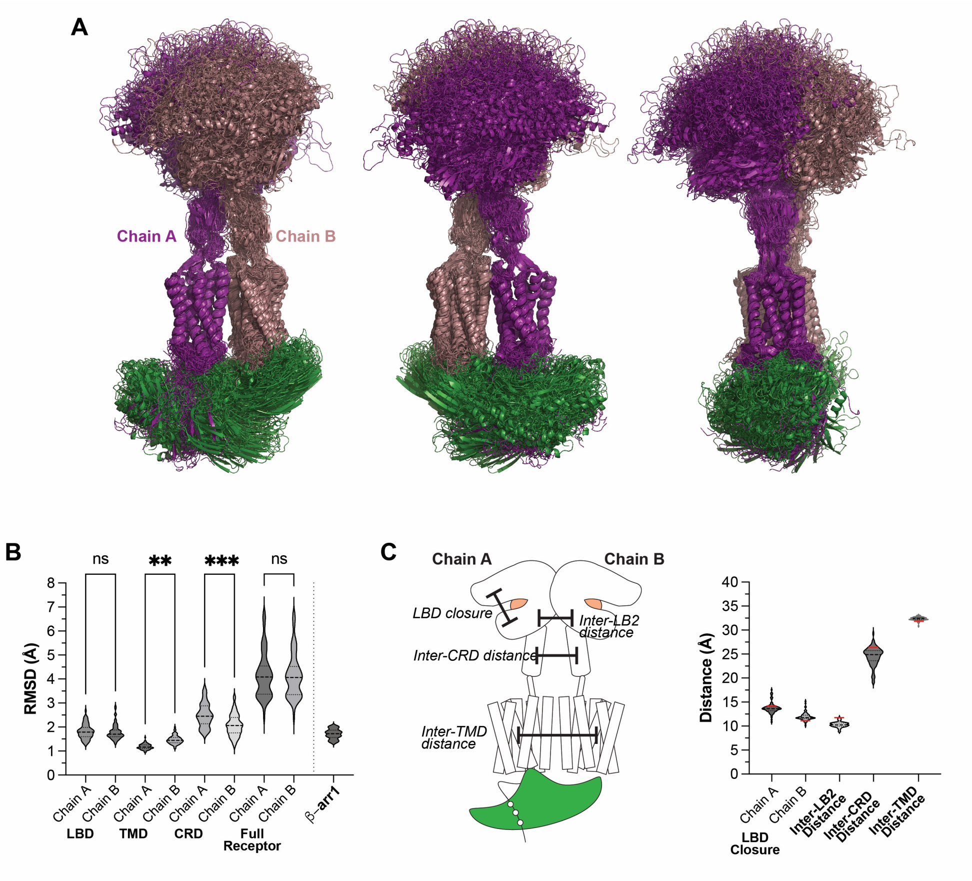


**Fig. S14. Analysis of mGluR8 conformation in MD simulations. (A)** Overlay of final frames of the 72 MD trajectories, aligned by the TMD (residues 580-850) of chain A of mGluR8. Overlay highlights the relative motion of both the LBD and β-arr1 to the TMD, as well as the lack of motion between the TMD of chain A and chain B of mGluR8. **(B)** Average root mean squared deviation (RMSD) of individual domains and chains of mGluR8 in each of the replicas are plotted as violin plots. We observe a statistical difference (1-way ANOVA with multiple comparisons; ** $p<0.01$; ***$p<0.001$) in RMSD of the chain A TMD and CRD compared to chain B, which may be due to β-arr1 binding to chain A (one way ANOVA). RMSD of β-arr1 is also small in all simulations (< 3Å) indicating β-arr1 is not experiencing large conformational changes. **(C)** Left, schematic highlighting various parameters measured in MD simulations to describe mGluR8 conformations. Right, distances plotted for each time point in all simulations. LBD closure is defined by the distance between residues 155 and 284, inter LB2 and CRD distances are the same as shown in Figure S13, between residues 229 and 553 in chain A and B, respectively. Inter TMD distance was calculated as the distance between the center of mass of the chain A and chain B TMD. Values from the starting structure are indicated with a red line.

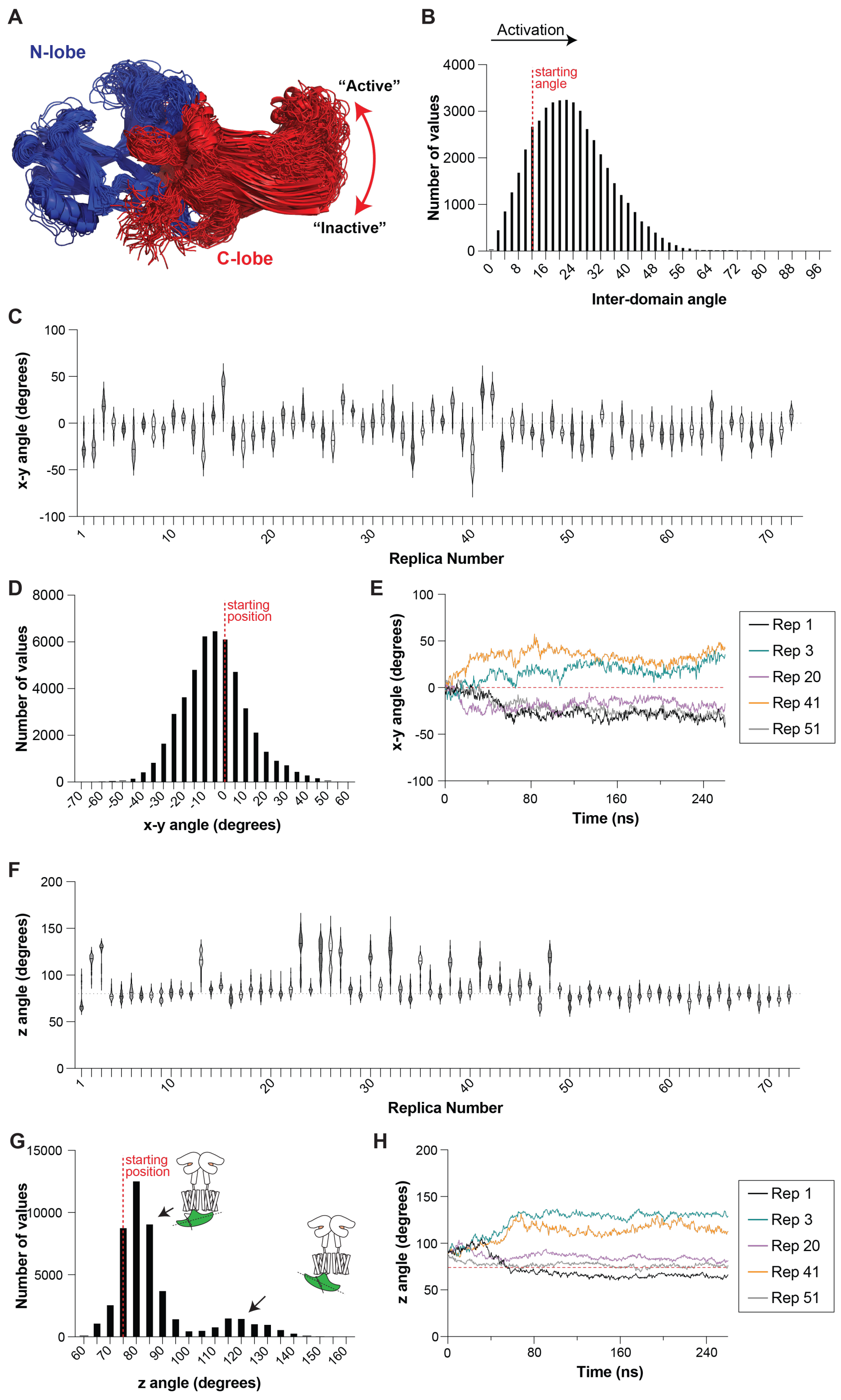


**Fig. S15. Analysis of β-arr1 conformation and orientation in MD simulations. (A)** Alignment of β-arr1 protomers from the final frame of each trajectory in the MD simulations, aligned by the N lobe (residues 1-175) showing the relative motion of the C lobe. **(B)** A histogram of the inter-domain angle of β-arr1, calculated by determining the angle between vectors derived from the principal axes of each β-arr1 lobe, is plotted as a proxy of β-arr1 activation. It has been previously shown that higher inter-domain angles are associated with β-arr1 activation. We find on average that simulations show an increase in β-arr1 inter-domain angle compared to the starting angle, though many β-arr1 conformations are populated throughout the trajectories. **(C)** X-Y angles for β-arr1 relative to mGluR8 are plotted for each timestep in each replica, shown as violin plots. **(D)** Cumulative histogram of x-y angles from all trajectories with the starting position of β-arr1 in our structural model highlighted in red. **(E)** Example traces for 5 replicas plotting β-arr1 x-y angle as a function of time. **(F)** β-arr1 z-angle plotted for each time point in each trajectory as violin plots. **(G)** Cumulative histogram of z angles of β-arr1 from all trajectories showing two populations of β-arr1 molecules defined as z greater or less than 100° with schematics highlighting the changes in β-arr1 position relative to mGluR8. **(H)** Representative traces of the same 5 trajectories as shown in (E) plotting z angle as a function of time.

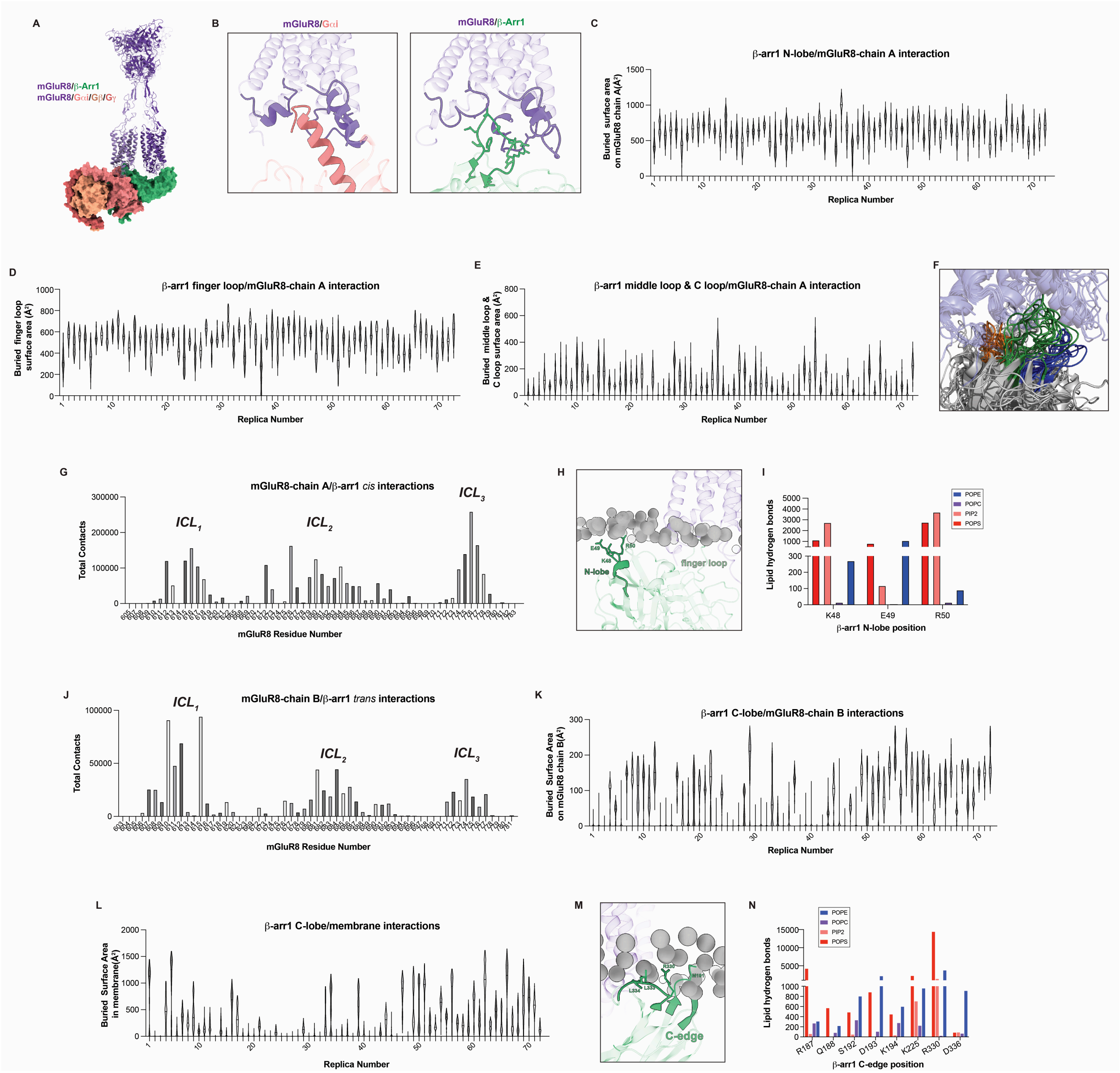


**Fig. S16. Analysis of mGluR8/β-arr1 interactions cryo-EM structures and MD simulations. (A-B)** Structural comparisons of mGluR8/β-arr1 model with G protein-bound mGluR8 structure showing distinct orientations (A) but highly similar interaction surface containing receptor intracellular loops (B). **(C)** Buried surface area between the β-arr1 N-lobe and mGluR8 chain A TMD for each time point in each trajectory, showing a consistent interaction surface that experiences some changes in the amount of buried surface as β-arr1 moves throughout the simulations. **(D-E)** Buried surface areas for β-arr1 finger loop (D), and middle and C loops (E) calculated as in (C). **(F)** Overlay of the final frames from a subset of trajectories shows interactions of β-arr1 loops with mGluR8 chain A TMD (finger loop: green, middle loop: blue, C loop: orange). **(G)** Quantification of contacts between mGluR8-chain A and β-arr1 across all simulations. **(H)** MD snapshot highlighting interaction between β-arr1 N-lobe with the membrane in a trajectory with a high z angle. **(I)** Lipid hydrogen bonds were calculated for β-arr1 residues highlighted in (G), showing a preference for interactions with negatively charged lipids. **(J)** Quantification of contacts between mGluR8-

chain B and β-arr1 across all simulations. **(K-L)** Buried surface area of the β-arr1 C-lobe with both mGluR8 chain B (K) and the membrane (L) highlight the variety of *trans* interactions observed by the β-arr1 C-edge in MD simulations. **(M)** Snapshot of β-arr1 C-edge embedding into plasma membrane. **(N)** Lipid hydrogen bonds are calculated for select β-arr1 C-edge residues, implicating arginine residues in driving charged, lipid-dependent membrane interactions.

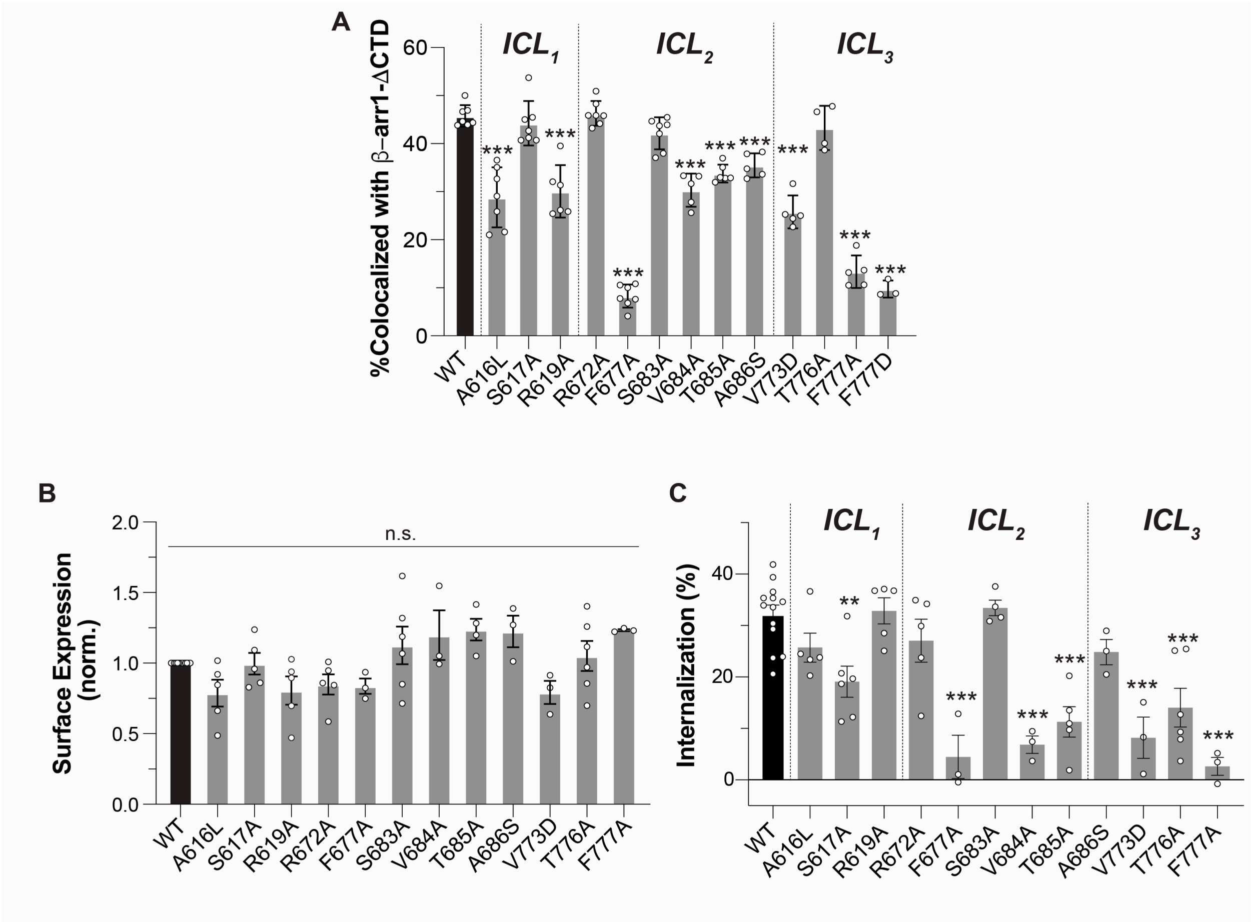


**Fig. S17. Further analysis of mGluR8 intracellular loop mutations. (A)** Quantification of degree of β-arr1-ΔCTD pulldown for mGluR8 core binding site mutations using SiMPull. **(B)** Surface expression analysis showing a lack of effect of mGluR8 core binding site mutations. **(C)** Quantification of degree of mGluR8 internalization for all core mutants showing a significant decrease relative to WT for many mutations. Points represent individual experiments. 1-way ANOVA with Dunnet's multiple comparisons test (to WT) is used in (A, B, C). **, p<0.01; ***, p<0.001. All data shown as mean ± SEM.

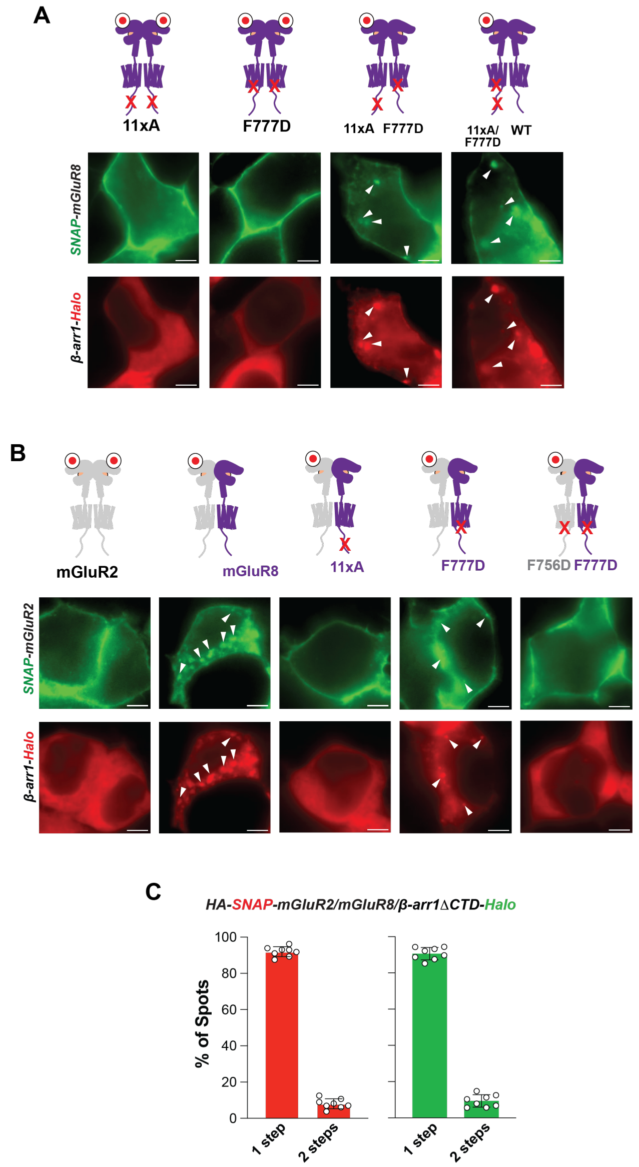


**Fig. S18. Further analysis of *cis* and *trans* mGluR8 and mGluR2/8 β-arr coupling. (A)** Representative images showing internalization and β-arr colocalization for a range of mGluR8 homodimer conditions. Arrows represent sites of receptor/β-arr1 intracellular co-localization. **(B)** Representative images showing internalization and β-arr colocalization for a range of mGluR2/8 heterodimer conditions. Arrows represent sites of receptor/β-arr1 intracellular co-localization. **(C)** Quantification of photobleaching step distribution supporting a 1:1:1 mGluR2:mGluR8:β-arr1 complex. Scale bar: 5 µm.

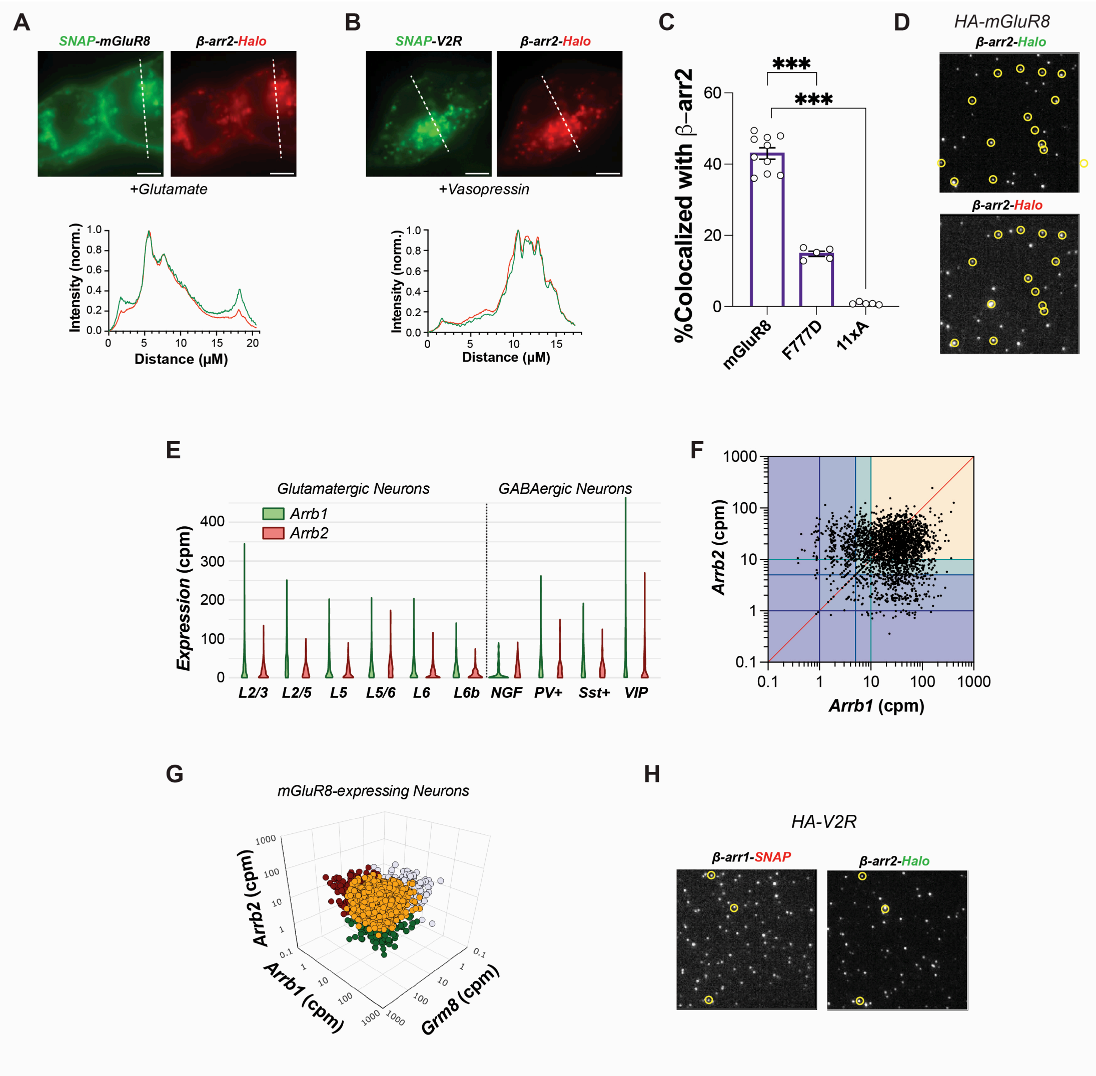


**Fig. S19. Further analysis of mGluR8/β-arr2 coupling. (A-B)** Representative images and line scans showing co-internalization of either mGluR8 (B) or V2R (C) with β-arr2. **(C)** Summary of β-arr2 SiMPull data for mGluR8 showing large effects of F777D and 11xA mutations. **(D)** Representative image showing colocalization of β-arr2-Halo labeled with two different fluorophores with HA-tagged mGluR8. **(E)**Violin plot showing Arrb1 (β-arr1) and Arrb2 (β-arr2) expression levels across neuronal classes. CPM=counts per million. **(F)** Quantification of RNA levels for *Arrb1* and *Arrb2* across neurons showing a wide range of ratios, including many cells which express high levels of both subtypes. Cutoffs of 1 (dark purple), 5 (blue) and 10 CPM (green) are depicted with the shadowed areas and corresponding division lines. **(G)** 3-dimensional scatter plot showing Arrb1, Arrb2, and Grm8 (mGluR8) expression levels across all Grm8-expressing cells **(H)** Representative image showing minimal colocalization between β-arr1-SNAP and β-arr2-Halo labeled with two different fluorophores with HA-tagged V2R. 1-way ANOVA with Dunnet's multiple comparisons test (to WT) is used in (C). ***, p<0.001. All data shown as mean ± SEM. Scale bar: 5 μm.

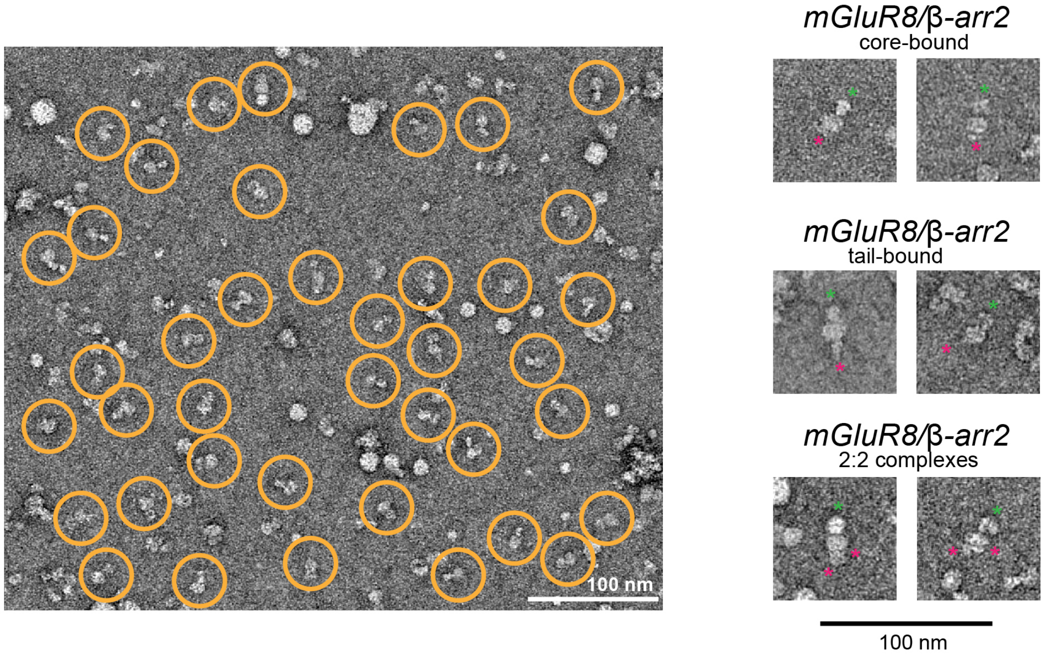


**Fig. S20. Representative mGluR8/β-arr2 negative stain particles.** EM micrograph showing identified mGluR8/β-arr2 complexes (orange circles), left, and representative single particles for each major class of interactions, right. Red stars highlight β-arr2 molecules and green stars highlight mGluR8 LBDs.

| | mGluR8 class 1 EMDB – xxx PDB - xxx | mGluR8 class 2 EMDB – xxx PDB - xxx | mGluR8/β-arr1 EMDB – xxx PDB - xxx | mGluR8/G protein class 1 EMDB – xxx PDB - xxx |
|---|---|---|---|---|
| **Data Collection** | | | | |
| Magnification | 105,000x | 105,000x | 105,000x | 105,000x |
| Voltage (kV) | 300 | 300 | 300 | 200 |
| Electron exposure (e–/Å$^2$) | 51.50 | 51.50 | 50.60 and 48.34 | 30.00 |
| Defocus range (μm) | 0.8-2.9 | 0.8-2.9 | 0.6-2.4 and 0.7-2.5 | 0.8-1.8 |
| Pixel size (Å) | 0.4125 (0.825 binned) | 0.4125 (0.825 binned) | 0.4125 (0.825 binned) | 1.16 |
| Symmetry imposed | C1 | C1 | C1 | C1 |
| Initial particle images (no.) | 3,400,746 | 3,400,746 | 9,511,601 | 5,988,809 |
| Final particle images (no.) | 338,052 | 298,012 | 189,406 | 142,834 |
| Map resolution (Å) | 3.1 | 3.1 | 2.9 | 5.9 |
| FSC Threshold | 0.143 | 0.143 | 0.143 | 0.143 |
| **Refinement** | | | | |
| Initial model used | AlphaFold2 multimer dimer of mGluR8 | AlphaFold2 multimer dimer of mGluR8 | mGluR8 alone class 1 | AlphaFold3 model of mGluR8/G protein |
| Model composition | | | | |
| Non-hydrogen atoms | 10547 | 10502 | 11104 | 17753 |
| Protein residues | 1564 | 1552 | 1587 | 2432 |
| Ligands | 4 | 4 | 3 | 3 |
| B factors (Å$^2$) | | | | |
| Protein | 125.52 | 116.61 | 147.19 | 117.52 |
| Ligand | 186.74 | 160.67 | 193.88 | 98.31 |
| R.m.s. deviations | | | | |
| Bond lengths (Å) | 0.002 | 0.004 | 0.003 | 0.005 |
| Bond angles (°) | 0.427 | 0.506 | 0.613 | 1.040 |
| Validation | | | | |
| MolProbity score | 1.98 | 2.56 | 1.94 | 2.24 |
| ClashScore | 8.40 | 14.31 | 10.47 | 17.66 |
| Poor rotamers (%) | 2.51 | 3.93 | 1.91 | 0.42 |
| Ramachandran plot | | | | |
| Favored (%) | 96.58 | 92.68 | 93.98 | 91.94 |
| Allowed (%) | 3.36 | 6.96 | 5.83 | 7.81 |
| Disallowed (%) | 0.06 | 0.07 | 0.32 | 0.25 |

**Table S1. Cryo-EM data collection, refinement, and validation statistics**. Information on cryo-EM data collection settings and refinement statistics for all mGluR8 models.

**Movie S1. Representative MD trajectories.** Six molecular dynamics trajectories are shown, with 5 of 6 matching the representative traces plotted in Fig. S15E and S15H. mGluR8 (chain A in blue, chain B in purple) and β-arrestin1 (green) are shown in cartoon representations and plasma membrane phosphate atoms are shown as orange spheres. Movies show full trajectories with 0.4ns time steps and highlight diversity in β-arrestin1 motion relative to mGluR8.
